# Nucleolus and centromere TSA-Seq reveals variable localization of heterochromatin in different cell types

**DOI:** 10.1101/2023.10.29.564613

**Authors:** Pradeep Kumar, Omid Gholamalamdari, Yang Zhang, Liguo Zhang, Anastassiia Vertii, Tom van Schaik, Daan Peric-Hupkes, Takayo Sasaki, David M. Gilbert, Bas van Steensel, Jian Ma, Paul D. Kaufman, Andrew S. Belmont

## Abstract

Genome differential positioning within interphase nuclei remains poorly explored. We extended and validated TSA-seq to map genomic regions near nucleoli and pericentric heterochromatin in four human cell lines. Our study confirmed that smaller chromosomes localize closer to nucleoli but further deconvolved this by revealing a preference for chromosome arms below 36-46 Mbp in length. We identified two lamina associated domain subsets through their differential nuclear lamina versus nucleolar positioning in different cell lines which showed distinctive patterns of DNA replication timing and gene expression across all cell lines. Unexpectedly, active, nuclear speckle-associated genomic regions were found near typically repressive nuclear compartments, which is attributable to the close proximity of nuclear speckles and nucleoli in some cell types, and association of centromeres with nuclear speckles in hESCs. Our study points to a more complex and variable nuclear genome organization than suggested by current models, as revealed by our TSA-seq methodology.

## Introduction

Early cytologists defined heterochromatin as chromosome regions which remained condensed post-mitosis throughout most of the interphase cell cycle [1–3]. They described heterochromatin as localizing preferentially at the nuclear and nucleolar peripheries across a wide range of species (reviewed in [4–8]). The advent of immunostaining and in situ hybridization methods indeed revealed varying preferential localization of centromeres, telomeres, as well as individual silenced genes to the nuclear and nucleolar peripheries [9–16]. Some silenced genes were also observed to preferentially localize adjacent to the pericentric heterochromatin (PCH) at the periphery of chromocenters, formed in species such as *Drosophila* and mouse through the clustering of centromeres [17–21]. However, the extent to which silenced genes and heterochromatin colocalize with PCH in species without prominent chromocenters, such as human, remains unclear. Importantly, most late-replicating heterochromatin regions were observed to localize at varying frequencies, higher than that observed for control euchromatic regions, among each of these different nuclear locations [22]. These microscopy observations led to the suggestion that the nuclear, nucleolar, and pericentric heterochromatin peripheries should be considered as forming one equivalent repressive compartment, with gene silencing and/or maintenance of this gene silencing correlated with localization to these nuclear compartments [23, 24].

The distinct biochemical composition of each of these nuclear compartments suggests different potential functional consequences when particular heterochromatin regions localizing to one or another of these different compartments. Additionally, localization of a given heterochromatic region to one or the another of these compartments would be expected to result in different nuclear localization of flanking euchromatic regions. These considerations raise questions of whether different types of heterochromatin might localize at different frequencies to these three compartments, the possible dynamics of this differential positioning throughout the cell cycle, physiological transitions, and cell differentiation, and the possible functional consequences of this differential positioning. A single genome-wide mapping approach that can measure and compare the relative localization of different heterochromatin regions to each of these three nuclear compartments would be highly beneficial for addressing these questions.

Previously, genome-wide, high-throughput sequencing methods which probe molecular proximity of the genome to particular proteins have been used as the most common approach to explore these questions. DamID and, more recently, ChIP-seq and Cut and Run or Cut and Tag molecular proximity methods have successfully mapped lamina associated domains (LADs) [5, 25–27]. A new “pA-DamID” method has overcome the limited time resolution of the original DamID method for LAD detection and has identified cell-cycle modulations of LAD associations [28], while single-cell DamID has measured contact frequencies of individual LADs [29]. DamID mapping of nucleolar associated domains using a nucleolar targeting peptide fused with the Dam methylase has been reported in K562 cells [30], and, more recently, using a nucleolar targeting peptide fused with histone H2B and Dam methylase, in mouse embryonic stem cells [31]. In other studies, Nucleolar Associated Domains (NADs) have been mapped by the sequencing of DNA co-purifying with isolated nucleoli [32–35]. These NADs largely overlap with LADs identified in the same cell lines, but also include H3K27me3-enriched inter-LAD domains. To date, PCH-associated domains (PADs) have only been mapped in mouse cells by 4C, using a major satellite DNA repeat as the viewpoint. While in somatic cells PADs largely overlap LADs and correspond to constitutively inactive genomic regions [36], unexpectedly in mouse ESCs a significant fraction of PADs instead overlapped with transcriptionally active, constitutive inter-LADs (iLADs) [36].

Recently a new genomic method, Tyramide Signal Amplification (TSA)-seq, was introduced which produces a signal proportional to distance rather than molecular proximity to the target protein. TSA-seq was able to map the genome relative to a nuclear compartment-nuclear speckles-which was unmappable using ChIP-seq [37]. TSA-seq also was able to map the genome relative to the nuclear lamina, in this case producing similar but complementary results to DamID [37]. For example, TSA-seq revealed that short inter-LADs (iLADs) still position near the nuclear lamina; conversely some LADs show an increased separation away from the nuclear lamina, perhaps interacting with nucleoli and/or the PCH [37]. Thus, TSA-seq either provides the ability to map the genome relative to nuclear locales that were previously unmappable or provides similar but complementary information to that provided by molecular proximity mapping methods.

We therefore developed and applied nucleolar and PCH TSA-seq to map and compare the relative association of heterochromatin with the nuclear lamina, nucleoli, and PCH across four different human cell lines: H1, HFF, HCT116, K562. We show varying association of different heterochromatin regions with different nuclear compartments both within and between cell lines.

## Results

### Extending TSA-Seq to measure cytological proximity to nucleoli and pericentric heterochromatin

We first considered which nucleolar and PCH subcompartments we should target, then identified suitable antibodies to mark these subcompartments, and finally optimized TSA staining conditions for each of the targeted subcompartments in HCT116 cells.

Certain markers from the Fibrillar Center (FC) and Dense Fibrillar Compartment (DFC) markers have the highest nucleolar to nucleoplasmic enrichment, which would minimize background staining. TSA labeling creates an exponential decay in tyramide free-radical concentration as a function of distance from the immunostaining target [37]. The interior localization of the FC and DFC within the nucleolus therefore would lower the tryamide free radical concentrations reaching the chromatin adjacent to the nucleolar periphery (Fig. S1a). Moreover, this exponential decay would result in little or even no signal for chromosome regions at the nucleolar periphery adjacent to the Granular Compartment (GC) but with no nearby FC. However, we reasoned that this reduced sensitivity using a FC or DFC target might also increase the spatial resolution of nucleolar TSA-seq mapping by labeling only regions closest to nucleoli. Conversely, antibodies targeting the GC would probe chromosome localization over a larger radius surrounding nucleoli. Following a similar logic, for marking the PCH we could choose either CENPB, which binds a subset of alpha-satellite repeats flanking centromeres [38], or CENPA, marking the centromeres themselves.

Screening anti-nucleolar antibodies, we found that whereas nucleophosmin (GC marker) imunostaining produced a strong nucleolar signal, particularly near the nucleolar periphery, it also produced a high nucleoplasmic background (Fig. S1b). Using the Human Protein Atlas (proteinatlas.org) [39] as a guide, we identified potential, low-background nucleolar markers including MKI67IP and DDX18, RNA polymerase I subunit E (Pol1RE) for the FC, and nucleolin for both the DFC and GC [40] (Fig. S1b). Antibodies tested for these markers produced low nucleoplasmic background and worked at high dilution (1:2000). Double labeling with nucleophosmin revealed MKI67IP distributed throughout the GC, whereas DDX18 localized in many small foci densely packed within the GC (SFig. 1b); therefore, both served as GC markers for TSA.

Both CENP-B and CENP-A immunostaining generated low nucleoplasmic background (Fig. S1b). CENP-B covered a larger fraction of the PCH whereas CENP-A decorates the actual, small dot-like centromeric region adjacent to the PCH; chromatin regions interacting with PCH could be some distance from the single, dot-like CENP-A staining on each interphase chromatid. Therefore, we chose CENP-B staining to probe genome interactions with the PCH.

Thus, we identified useful antibodies for TSA staining of the PCH and different nucleolar compartments which produced both low background and strong TSA signals (Fig. 1A).

**Figure 1:**
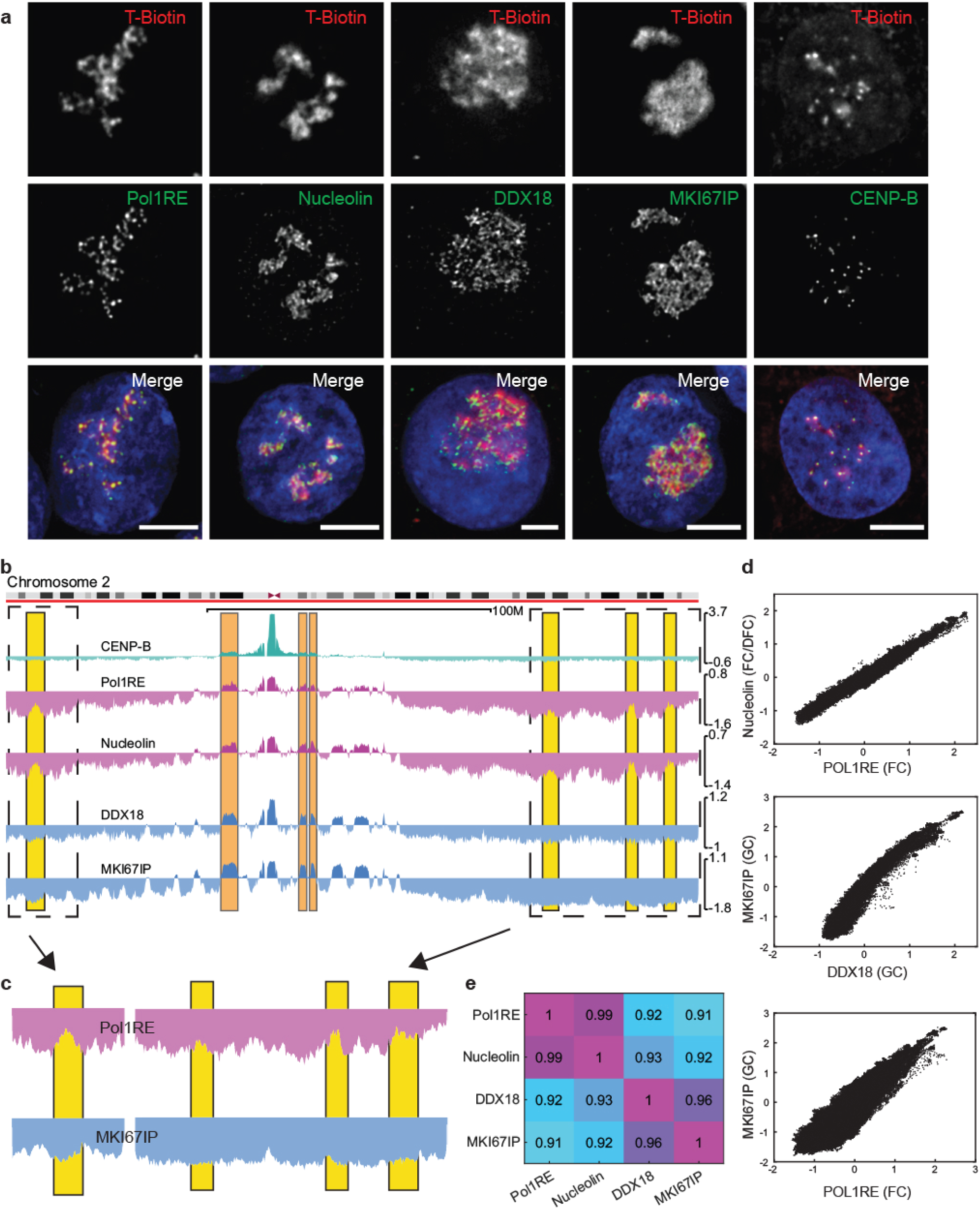
Identification and comparison of molecular markers for nucleolar and pericentric heterochromatin (PCH) TSA-seq: **a.** Immunostaining (green) and TSA (red, streptavidin labeling of tyramide biotin) examples using nucleolar markers POL1RE (FC), Nucleolin (DFC/FC), DDX18 and MKI67IP localized here to the GC, and CENP-B as a pericentric heterochromatin marker. DNA staining (blue); Scale bar = 5 um. **b&c.** TSA-seq Chr2 Nucleome Browser views produced in HCT116 cells against markers (from top to bottom) CENP-B, POL1RE, Nucleolin, DDX18, and MKI67IP. Orange highlights-chromosome regions appearing as peaks in both CENP-B and Nucleolar TSA-Seq; yellow highlights-chromosome regions appearing as peaks within valleys in FC/DFC TSA-Seq but valleys in GC TSA-Seq. **d.** High correlation between TSA-seq produced using different nucleolar markers: 2D Scatterplots (top to bottom)-Nucleolin (DFC/FC) vs Pol1RE (FC), MKI67IP (GC) vs DDX18 (GC), MKI67IP (GC) vs POL1RE (FC) TSA-Seq. **e.** Pearson correlation heat map of TSA-Seq for different nucleolar markers.

We next surveyed different TSA-seq 2.0 staining conditions [41] to optimize the TSA genomic labeling. Using intermediate tyramide-biotin concentrations and staining times (“Condition C” from our previous study) [41], all four nucleolar markers produced similar patterns of TSA-Seq scores (log_2_(pulldown/input) appropriately normalized by read count [37]) (Fig. 1). Notably, NOR-adjacent chromosome regions displayed the highest TSA-seq scores (Fig. 2, left ends of Chr 13,14,15, 21, 22) but centromeres of metacentric chromosomes generally also contained strong TSA-seq peaks (Fig. 1b, Fig. 2). CENP-B TSA-Seq detected major peaks overlapping the centromere and smaller “satellite” peaks in regions flanking the centromere which typically overlap with nucleolar TSA-seq peaks (Fig. 1b, orange highlights). We hypothesize that the nucleolar association of centromeres contributes to the correlation of these PCH TSA-seq satellite peaks with nucleolar TSA-seq peaks.

**Figure 2:**
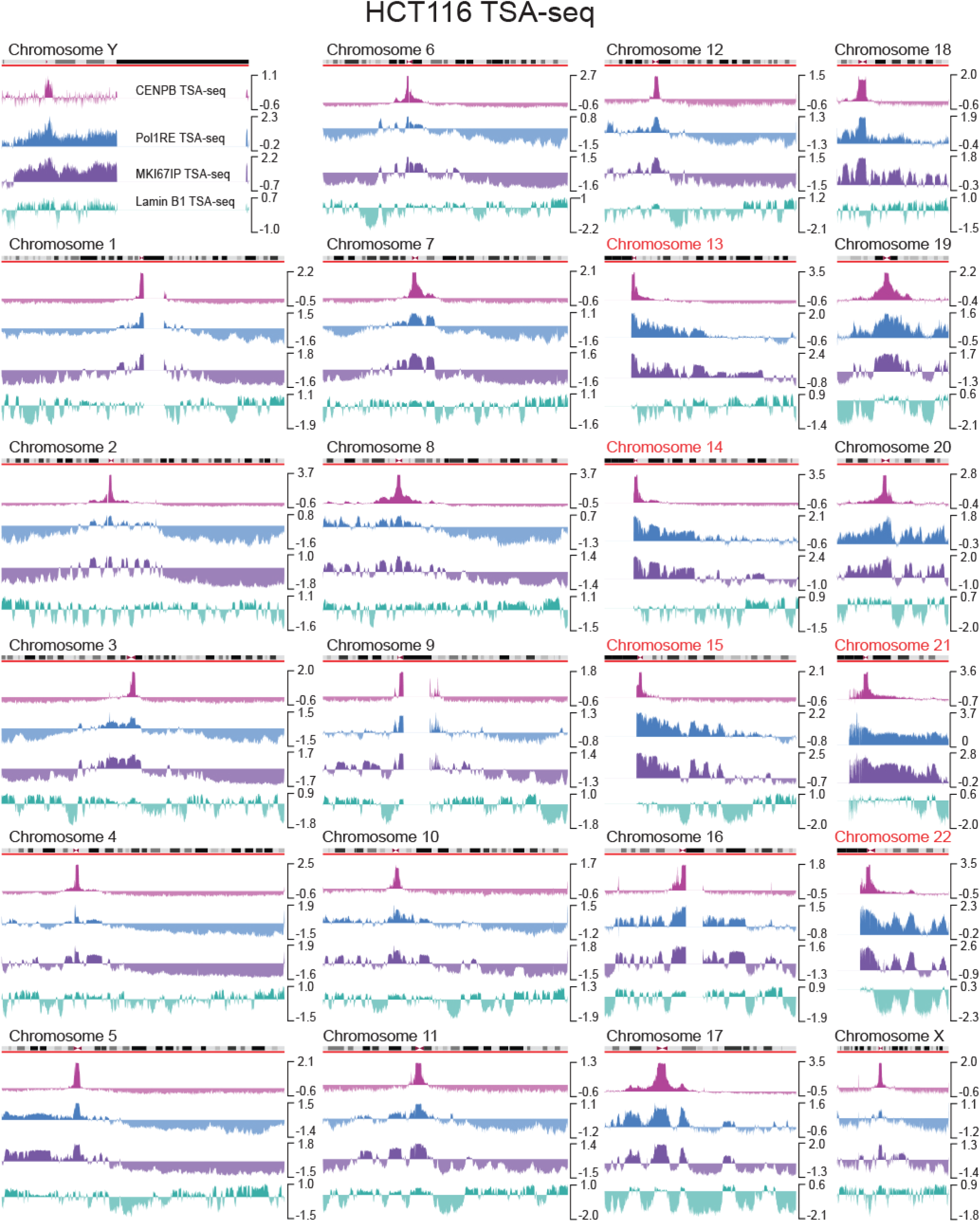
Comparison of CENP-B, POL1RE, MKI67IP, and Lamin B1 TSA-seq across all chromosomes in HCT116 cells. Nucleome Browser views are based on Hg38 assembly. NOR containing chromosomes are highlighted in red.

Closer examination, however, reveals small differences between the FC/DFC (nucleolin, Pol1RE) versus GC (MKI67IP, DDX18) target TSA-seq. For example, “peaks-within-valleys” in the FC/DFC TSA-seq do not align with similar features in the GC TSA-seq (Fig. 1b&c, yellow highlights). Moreover, TSA-seq scatterplot comparisons showed the strongest correlations between data sets produced using antibodies targeting the same or spatially similar nucleolar compartments (Fig. 1d&e).

Moving forward, we choose MKI67IP and Pol1RE as markers for the GC and FC compartments, respectively, due to their higher TSA-seq dynamic range (Fig. 1b). We also enhanced TSA-labeling by using an improved protocol (TSA-seq 2.0 Condition E) which reduced the required cell numbers [41]. Compared to data obtained using the Condition C protocol, the increased TSA labeling obtained using Condition E produced similar TSA-seq results for MKI67IP, Pol1RE, and CENP-B but with slightly reduced dynamic range (Fig. S1c-d) [41].

We then performed TSA-seq to produce replicate datasets for MKI67IP, Pol1RE, CENP-B as well as LMNB1 in all four cell lines analyzed-HCT116, H1, K562, and hTERT-HFF. This data complemented previously generated SON TSA-seq in all four cell lines [41].

### Specificity of nucleolar and PCH TSA-seq supported by initial examination of genome-wide maps

In all cell types analyzed, we observed maximal CENP-B TSA-seq signals over chromosome centromeric regions, signal decay away from centromeric regions, and a small number of smaller peaks flanking centromeric regions, as we had seen in the HCT116 cells. Together, these data validate the specificity of the CENP-B TSA-seq (Fig. 1b, Fig. S2, Fig. 2).

Inspection of the HCT116 nucleolar TSA-seq (Fig. 2) reveals consistency with several known features of nucleolar chromosome association. The maximum nucleolar TSA-seq signals for all four markers appear in the hg38 chromosome assembly over regions on the NOR-containing chromosomes (13,14,15,21,22) immediately adjacent to their repeat-rich p-arms containing the ribosomal gene repeats; the p-arms and NORs within these p-arms are missing from this assembly (Fig. 2). Remapping to the newer T-to-T complete human chromosome assembly [42] revealed the highest nucleolar TSA-seq signals over the NOR regions themselves, with the second highest nucleolar TSA-seq signals over the non-NOR p-arm regions known to associate with the nucleolar periphery (Fig. 3a) [43].

**Figure 3:**
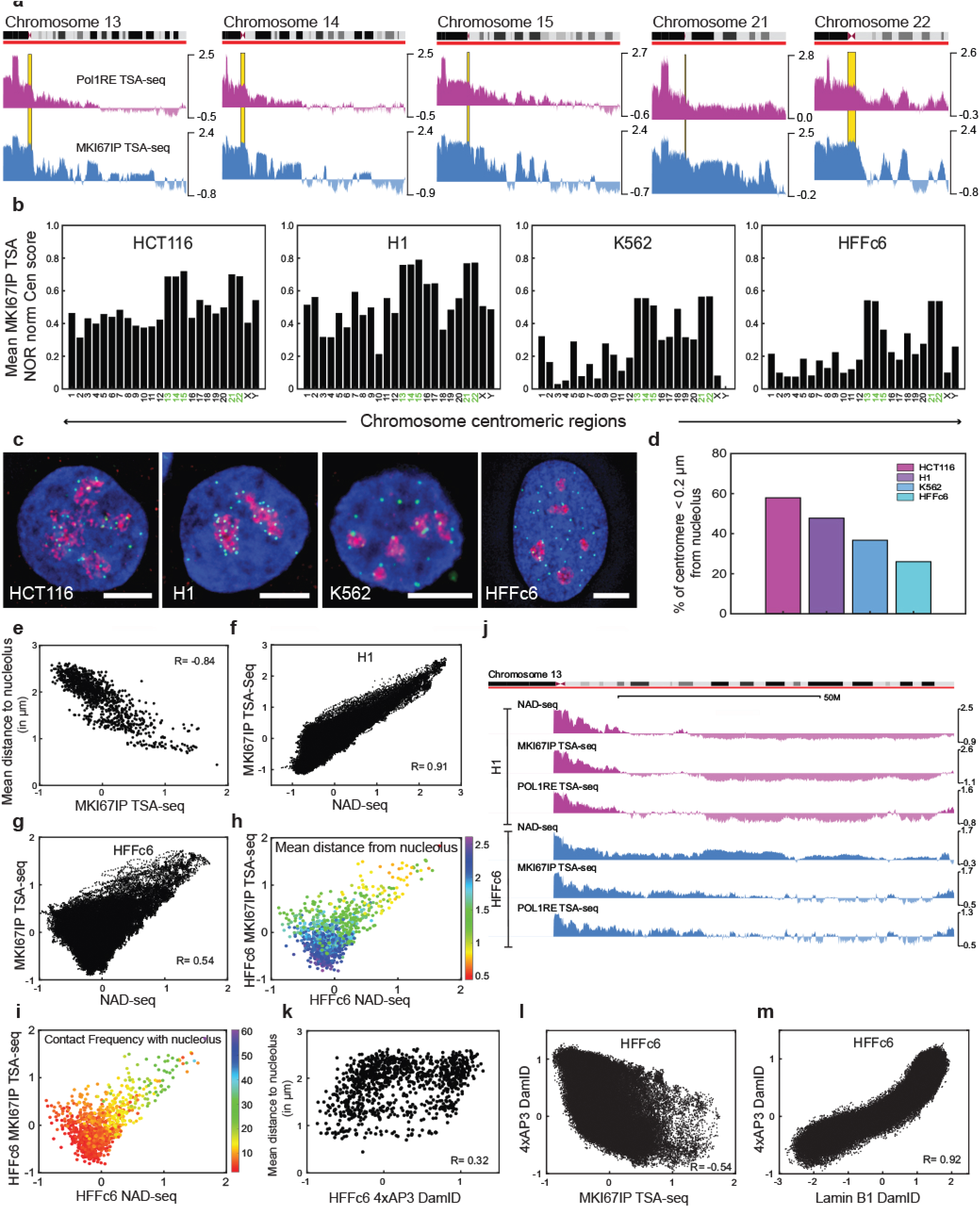
Validation of nucleolar TSA-seq. **a** TtoT genome assembly reveals highest nucleolar HCT116 TSA-seq signals over p-arms containing NORs and rDNA repeats. Browser views of POL1RE (top) and MKI67IP (bottom) TSA-Seq for NOR-containing chromosomes (left to right) Chr13, Chr14, Chr15, Chr21, Chr 22. Yellow highlights show centromere positions. **b-d**. Variation in nucleolar TSA-seq signals across different cell lines over centromere regions matches variable centromere localization to nucleoli seen by microscopy. Mean MKI67IP TSA-seq scores (y-axis) over centromere regions (x-axis) (b) shows highest values for NOR-containing chromosomes in general. Non-NOR containing chromosomes show higher and near constant centromere nucleolar TSA-seq signals in HCT116 and H1 cells but lower and more variable signals in HFF and K562 cells. Immunostaining (c) shows a high percentage (d, y-axis) of centromeres (CENP-A, green) associated with nucleoli (MKI67IP, red) (in red) in HCT116 and H1 but lower percentage (d) in K562 and HFF (DNA (DAPI) is blue; scale bars = 5 um); bar plots (d) showing colocalization percentages are from measurements from 50 nuclei for each cell type (d, x axis), using a distance threshold of less than 0.2 um. **e-i**. Nucleolar TSA-seq correlates both with fibroblast nucleolar distance measurements and NAD-seq. Scatterplots of distance to nucleolus (FC) (IMR90) versus MKI67IP TSA-seq (HFF) (e), MKI67IP TSA-seq versus NAD-seq in H1 (f) and HFF (g), MKI67IP TSA-seq versus NAD-seq in HFF with either distance to (h) or contact frequency with (i) to nucleoli (IMR90) superimposed in color-coding. **j**. Browser views comparing MKI67IP (GC) and POL1RE (FC) TSA-seq with NAD-seq in H1 (top tracks, purple) and HFF (bottom tracks, blue). **k-m**. Nucleolar DamID correlates poorly with microscopy distance measurements and nucleolar TSA-seq but correlates instead with lamin B1 DamID. 2D scatter plots of distance to nucleoli (IMR90) versus 4xAP3 nucleolar DamID (HFF) (k), 4xAP3 nucleolar DamID versus MKI67IP TSA-seq in HFF (l) or lamin B1 DamID (m).

To compare centromere associations with nucleoli across and within cell types, we normalized centromere MKI67IP TSA-seq scores to the maximum nucleolar TSA-seq scores of NORs in the T-to-T genome assembly. In HCT116, the second highest MKI67IP TSA-seq peaks are located at or immediately adjacent to centromeres (Fig. 2). Centromeres of non-NOR containing chromosomes showed near uniform MKI67IP TSA-seq scores of ~40-50% of the TSA-seq scores over the NORs while centromeres of the NOR-containing chromosomes showed ~70% the NOR scores (Fig. 3b). Centromere nucleolar TSA-seq scores were similar in H1 cells but centromeres of non-NOR containing chromosomes showed both substantially reduced and more variable nucleolar TSA-seq scores in HFF and K562 cells (10-30% in HFF, 5-30% in K562) (Fig. 3b). Variations in MKI67IP TSA-seq centromere scores across the four cell lines paralleled variations in frequencies of centromere/nucleoli colocalization measured by light microscopy (Fig. 3c-d) [44].

### Genome-wide validation of nucleolar TSA-seq

Other nucleolar TSA-seq features were unanticipated, raising validation concerns. Specifically, some previous NAD-seq studies measured similar levels of nucleolar association across most LADs [35], as did nucleolar DamID produced using Dam methylase fused to a nucleolar targeting peptide [30]. In contrast, although MKI67IP TSA-seq scores in HCT116 are higher over LADs than flanking iLADs, they change from peaks to “peaks-within-valleys” or flat valleys moving towards the ends of long metacentric chromosome arms (Fig. 2). In other cell types, many LADs even appear as MKI67IP TSA-seq local minima rather than local maxima, while small peaks and “peak-within-valley” local maxima of MKI67IP TSA-seq align instead with iLAD subregions corresponding to nuclear speckle (SON) TSA-seq peaks, which correlate with gene dense, highly transcriptionally active genome regions [37, 41].

For validation purposes, we therefore compared nucleolar TSA-seq with two orthogonal measurements of genome association with nucleoli. First, we analyzed highly multiplexed, genome-wide, immuno-FISH imaging of chromosomal nucleolar association [45] (Fig. 3, Fig. S3). Strongly validating the TSA-seq from HFF human fibroblasts, both the MIKI67IP (Fig. 3e, 0.84 Pearson correlation coefficient) and POL1RE (Fig. S3b, 0.80 Pearson correlation coefficient) TSA-seq scores correlated inversely with the measured distances of ~1000 FISH probes spaced evenly across the genome to the nearest nucleolar DFC in IMR90 human fibroblasts.

As a second genome-wide validation test (Fig. 3f-i), we compared nucleolar TSA-seq with NAD-seq, in which nucleolar associations were measured by deep sequencing of DNA associated with biochemically purified nucleoli. In previous analyses, NAD peaks were identified as peaks in the local signal “contrast”, computed by subtracting the background signal defined by the mean of a sliding window [34, 46]. Such background subtraction artificially enhances the correlation between LADs and NADS. Instead, without this background subtraction, nucleolar TSA-seq and NAD-seq signals are highly correlated in H1 cells, as appreciated in both genome browser views (Fig. 3j, Fig. S3a) and scatterplot analysis (Fig. 3f, Fig. S3c). In HFF, the nucleolar TSA-seq versus NAD-seq correlation is significant but not as strong (Pearson Correlation coefficients 0.91 versus 0.54 (MKI67IP), 0.71 versus 0.41 (FC)) (Fig. 3g, Fig. S3d).

Whereas nucleolar TSA-seq reports on distance from nucleoli, NAD-seq measures the fraction of loci that maintain contact through the biochemical fractionation of nucleoli. Therefore, we expect greater deviations between TSA-seq and NAD-seq over genomic regions with lower nucleolar contact frequencies and higher variation in nucleolar distances. Indeed, genomic regions showing high MKI67IP TSA-seq show higher correlation with NAD-seq; these regions include smaller chromosomes, NOR-containing chromosomes, NORs and chromosome regions flanking NORs, and near centromeres in non-NOR containing chromosomes (Fig. 3f, Fig. S3A, and data not shown). Conversely, chromosome regions with lower nucleolar association show larger deviations between the TSA-seq versus NAD-seq values, particularly in HFF versus H1 cells. This is appreciated first by the widening of the off-diagonal scatterplot bins at lower values of both nucleolar TSA-seq and NAD-seq (Fig. 3g, Fig. S3d) and second by the superimposing of mean nucleolar distances (Fig. 3h) and nucleolar contact frequencies (Fig. 3i) onto these scatterplots. Nucleolar contact frequencies vary for loci with similar mean distances, contributing to the decreased correlation at lower values of both TSA-seq and NAD-seq.

In contrast, nucleolar DamID [30] in HFF did not correlate well with either mean distance to nucleoli in IMR90 (Fig. 3k), NAD-seq (Fig. S3E), or MKI67IP TSA-seq (Fig. 3l). Instead, for unknown reasons, nucleolar DamID strongly correlated with LMNB1 DamID (Fig. 3m) and LMNB1 TSA-seq (Fig. S3F).

Based on these genome-wide FISH and NAD-seq comparisons, we conclude that nucleolar TSA-seq measurements produce valid measurements of mean distance to nucleoli.

### Shorter chromosome arms show increased nucleolar association

Chromosome paint FISH previously demonstrated a more interior location for human NOR-containing chromosomes as well as smaller chromosomes 16-20 [47], possibly due to an increased association of these chromosomes with nucleoli. MKI67IP TSA-seq in both HCT116 (Fig. 4a) and HFF (Fig. S4a) cells shows a general trend of increased nucleolar association with decreasing chromosome lengths, but with some deviations (e.g., Chr 6-12, 17, 19, X, Fig. 4a).

**Figure 4:**
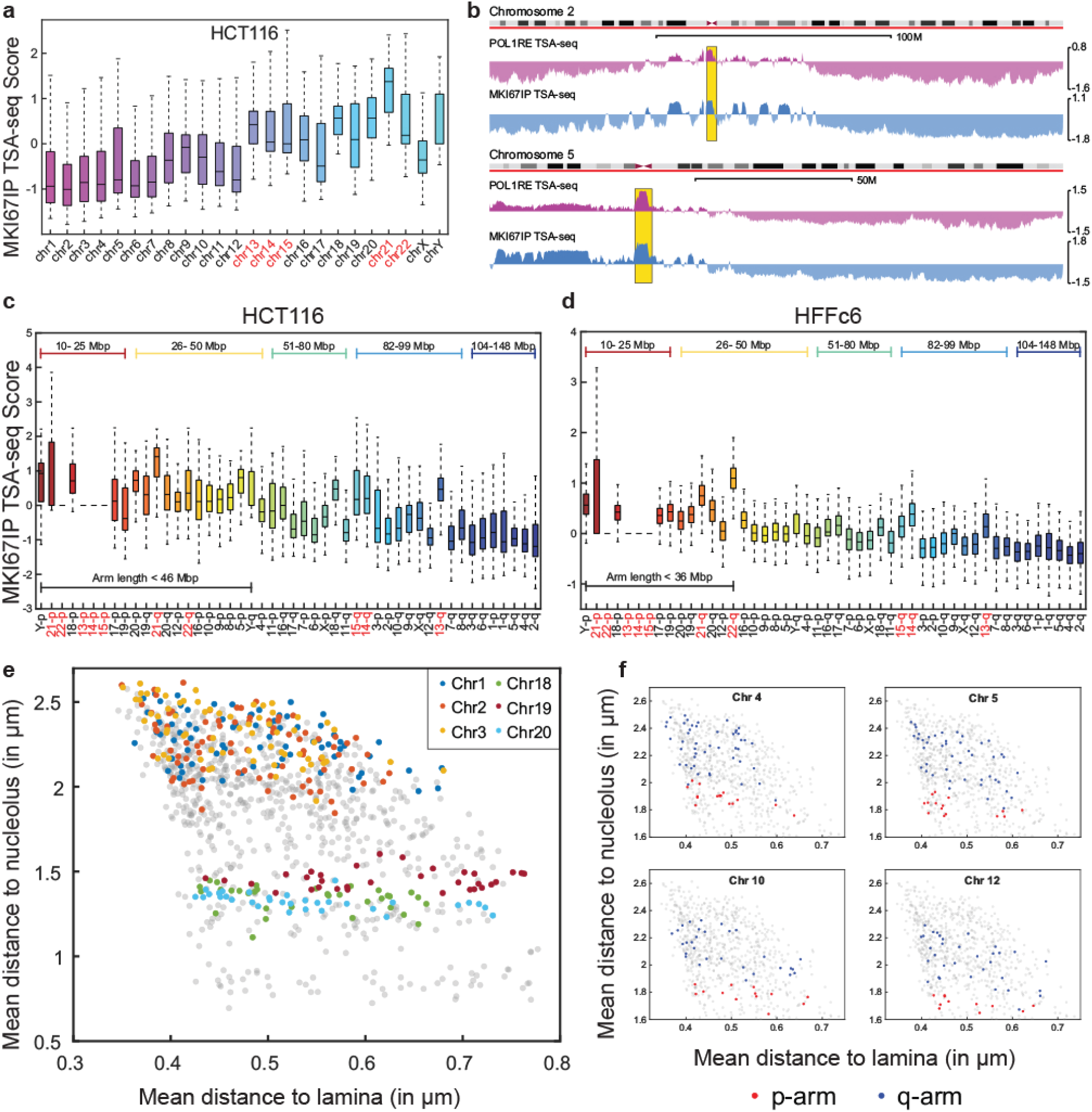
Increased nucleolar association with decreasing chromosome size deconvolved into increased nucleolar association of chromosome arms below critical length. **A.** NOR containing chromosomes (highlighted in red, x-axis) and smaller chromosomes (Chr 16-20) have higher MKI67IP TSA-Seq scores (y axis) compared to larger chromosomes (Chr 1-12) in HCT116 cells. **b.** Browser views of POL1RE (FC) and MKI67IP (GC) TSA-Seq in HCT116 show increased nucleolus TSA-Seq scores on the smaller p-arm of Chr 5 (bottom) compared to p arm of Chr 2 (top). **c-d.** Chromosome arm lengths less than 46 Mbp in HCT116 (c) and 36 Mbp in HFFc6 (d) show higher MKI67IP TSA-Seq scores (y axis). **e.** Scatterplots of mean distance to nucleolus (y-axis) versus mean distance to lamina (x-axis) for genomic regions on larger (Chr1-3) versus smaller (Chr18-20) chromosomes (Chr1-3). **f.** Four chromosome scatterplots of mean distance to nucleolus (y-axis) versus mean distance to lamina (x-axis) show p-arm genomic regions (red dots are closer to nucleoli than q-arm genomic regions (blue dots).

However, MKI67IP TSA-seq reveals an even stronger but nonlinear correlation between chromosome arm length and nucleolar association (Fig. 4, Fig. S4). For example, the short Chr 5 p-arm shows a pronounced increased nucleolar association along its length as compared to the steady drop in nucleolar TSA-seq moving distal from the centromere across the moderate length p-arm of Chr2 (Fig. 4b). Ignoring NOR-containing arms and the heterochromatic Y p-arm, chromosomes arms shorter than ~46 Mbp in HCT116 show generally higher nucleolar TSA-seq scores (Fig. 4c), that are relatively constant with arm length (Fig. 2). Exceptions include the q arms of the NOR chromosomes, as well as the q-arm of chromosome 18, which deviate towards higher nucleolar TSA-seq scores. The former is expected due to the near 100% association of NOR-containing p-arms with the nucleolus. The entire chromosome 18 territory previously had been shown to be localized towards the nuclear periphery in rounder but not flat cells [47, 48].

A similar trend between nucleolar association and arm length is seen in HFF, K562, and H1 except that the size beyond which nucleolar association decreases changes from ~46 Mbp to ~36 Mbp and the anomalous elevation in the chromosome 18 q-arm is lost in H1 (Fig. 4d, Fig. 4Sb-c). Mining the IMR90 multiplexed immuno-FISH data set [45] supports both the decreased distance from nucleoli of probes located on shorter (Chr18-20) versus longer chromosomes (Chr1-3) (Fig. 4e) and the variation in mean nucleolar distances as a function of chromosome arm length (Fig. 4f) inferred from the MKI67IP TSA-seq.

### Most LADs/NADs associate with both the nuclear lamina and nucleolar periphery but with different frequencies

Having extensively validated the nucleolar TSA-seq, we then used this method to determine whether different heterochromatin regions associate at the same or different relative frequencies with the nuclear lamina versus the nucleolar periphery using a single, and therefore more readily comparable, methodology.

Comparing HCT116 nucleolar versus nuclear lamina TSA-seq, we noted large differences in the relative distribution of many LADs relative to the nucleolus versus nuclear lamina (Fig. 2). For example, large LMNB1 TSA-seq peaks can correspond to very different nucleolar TSA-seq signals ranging from large to small peaks, a peak-within-valley local maxima, or a deep valley in the nucleolar TSA-seq signal. Similarly, many large nucleolar TSA-seq peaks show noticeably smaller LMNB1 peaks than typical for flanking LAD regions which have lower nucleolar TSA-seq scores (e.g. middle regions of Chr2, Chr3). A second overall trend is that the relative strengths of LMB1 TSA-seq peaks tend to increase near the ends of long chromosome arms while nucleolar TSA-seq signals decrease (Fig. 2).

We conclude that different LAD regions show varying relative association frequencies with the nuclear lamina versus nucleolar peripheries, with most LADs showing some level of increased affinity to both compartments.

### Identification of two distinct subsets of LADs/NADs localizing differentially between the nuclear periphery and nucleolus in different cell lines

LADs typically have been conceptualized and analyzed as a set of heterochromatin domains with similar biochemical and functional properties [5]. However, an analysis of an ~1 Mbp LAD region flanking the *HBB* locus, suggested at least three different nuclear lamina targeting mechanisms each operating within different regions of this LAD [49]. To ask whether different LAD “flavors” might be recognized by their varying association with different nuclear compartments, we focused on LADs which change nuclear positioning across different cell types, rationalizing that this would avoid the confounding bias of chromosome arm position on relative nuclear lamina versus nucleolar positioning.

Comparing the relative localization of LADs relative to the nuclear lamina versus nucleoli across the four cell lines identified two LAD subsets showing opposite biochemical and functional properties from each other (Fig. 5, Fig. S5, Table S1). Subset1 LADs are more repressed and later replicating while subset2 LADs are less repressed and earlier replicating than the set of all LADs. Although both LAD subsets were identified by their differential nuclear localization between different cell lines, their distinctive biochemical and functional properties are similar across all cell lines.

**Figure 5:**
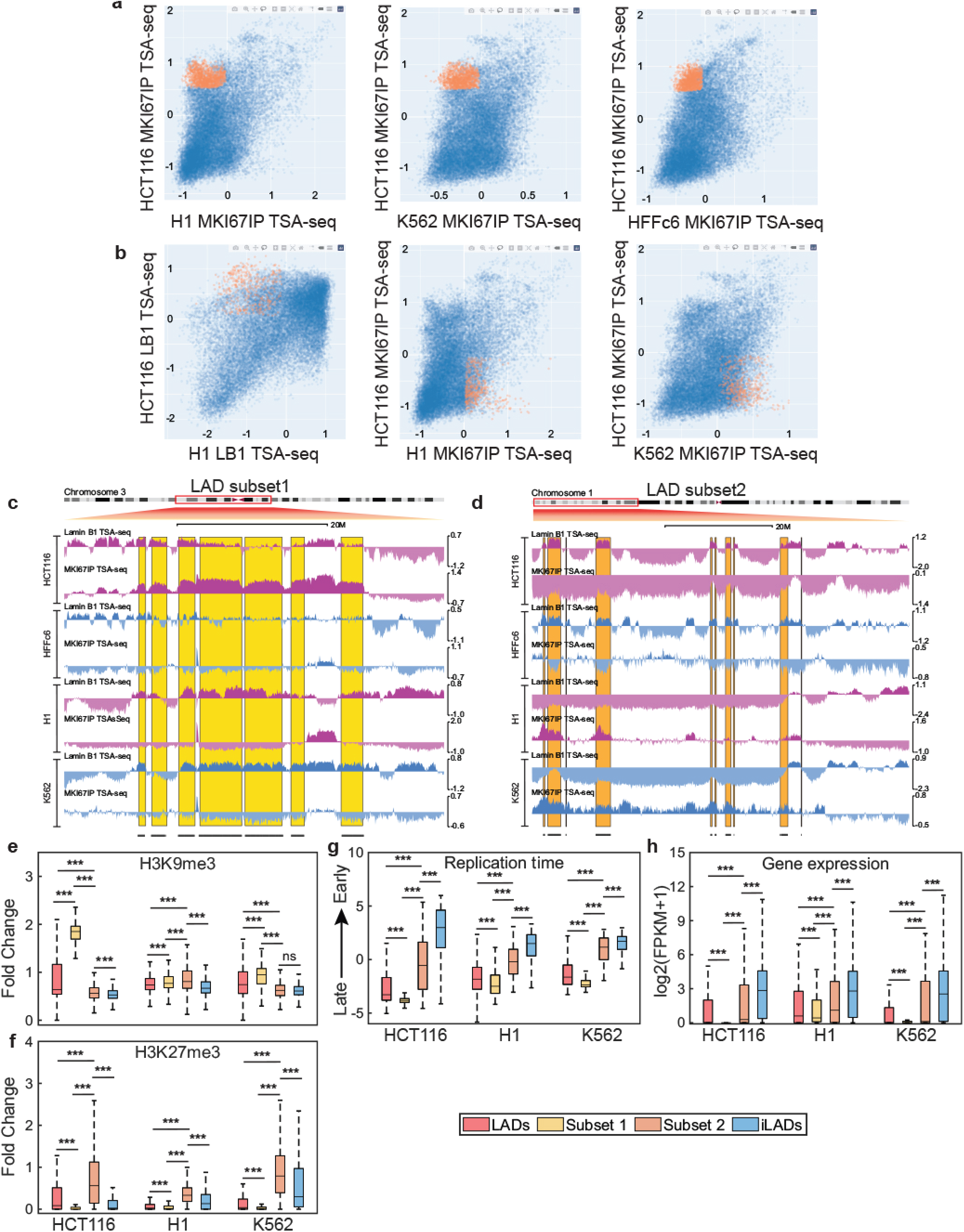
LAD subsets identified based on their cell-type specific nuclear localization show distinctive histone marks, gene expression levels, and DNA replication timing. **a&b.** Scatter plot tool was used to simultaneously select LAD subset1 genomic bins (highlighted in orange) which localize close to the nucleolus specifically in HCT116 cells (high MKI67IP TSA-Seq in HCT116, low in HFF, K562) (a) or LAD subset2 genomic bins which switch from close to the nucleolus in K562 and H1 to close to the nuclear lamina in HCT116 (high lamin B1 TSA-Seq in HCT116, low in lamin B1 TSA-Seq in K562 and H1 and high MKI67IP TSA-Seq in K562 and H1 but low in MKI67IP TSA-Seq in HCT116) (b). **c&d.** Chr3 (c) or Chr1 (d) browser views of Lamin B1 TSA-Seq and MKI67IP TSA-Seq in HCT116, HFFc6, H1 and K562 and segmented bins (bottom track, lines). LAD subset1 (c) and LAD subset2 regions (d) are highlighted in yellow and orange respectively. **e-f.** Relative to all LADs, LAD subset1 are enriched in H3K9me3 (e), low in H3K27me3 (f), replicate late in S-Phase (g) and have lower gene expression (h) while LAD subset2 LADs are enriched in H3K27me3 (f), low in H3K9me3 (e), replicate middle in S-phase (g) and have intermediate gene expression (h).

### LAD subset1

LAD subset1 was recognized by their conversion from low nucleolar association in three cell lines (H1, K562, HFF) to a relatively high nucleolar association, combined with a noticeable reduction in lamin B1 TSA-seq scores, in HCT116 cells (Fig. 5a&c). For analysis, we defined specific windows for MKI67IP and lamin B1 TSA-Seq scores (Fig. S5a-b) which identified such LADs (Fig. 5c) (see Materials and Methods).

Biochemically, LADs subset1 is distinguished most notably by higher H3K9me3-enrichment as compared to the set of all LADs. This enrichment is present in all four cell lines (Fig. 5e, HCT116, H1, K562; Fig. S5c, HFF) but is exceptionally elevated in HCT116 cells. In contrast, levels of H3K27me3 are low relative to the set of all LADs (Fig. 5f). Conversely, histone marks associated with active chromatin-H3K27ac (Fig. S5d, HFF; Fig. S5h, HCT116, K562, H1), and both H3K9ac and H3K4me1 (Fig. S5g&i, HCT116, H1, K562)-are noticeably depleted in LADs subset1 relative to the set of all LADs in each of the three or four cell lines where this data is available. Curiously, the active H3K4me3 mark instead is significantly elevated relative to the sets of all LADs and all iLADs specifically in LADs subset1 in HCT116; instead, this active mark is depleted in LAD subset1 in H1 and K562 (Fig. S5j).

Functionally, LADs subset1 shows significantly later DNA replication timing than the set of all LADs in all four cell lines, as assayed by 2-fraction Repli-seq (Fig. 5g, Fig. S5e), while average gene expression levels are significantly lower relative to all LADs in all four cell lines (Fig. 5h, Fig. S5f).

### LAD subset2

LAD subset2 was recognized by their conversion between cell types from high lamin B1 and low MIKI67IP TSA-seq scores to low lamin B1 and elevated MKI67 IP TSA-seq scores (Fig. 5b,d). Browser views revealed subset2 LADs localize predominately towards the ends of long chromosome arms and are largely LADs in HFF and HCT116 cells versus nucleolar-associated iLADs in K562 and H1 cells.

For analysis, we defined appropriate windows for MKI67IP and lamin B1 TSA-Seq scores (see Materials and Methods) which effectively identify those same regions (Fig. 5d, black bars in bottom browser track). Fig. 5d shows an ~60 Mbp region containing several subset2 LADs on Chr1 which alternates between nuclear speckle (SON) and lamin B1 TSA-peaks in HCT116 and HFF cells versus MKI67IP TSA-seq peaks in H1 and K562 cells (Fig. 5d, orange shaded regions).

LAD subset2 is more highly enriched in H3K27me3 than either the sets of all LADs or all iLADs in HCT116, H1, and K562 where data is available (Fig. 5f). LAD subset2 H3K9me3 is lower in HCT116 and K562 but higher in H1 and HFF cells relative to the set of all LADs (Fig. 5e, Fig. S5c). Active histone marks H3K27ac, H3K9ac, H3K4me1, H3K4me3, are all elevated in LAD subset2 relative to all LADs (Fig. S5d, g-j). In H1 hESCs, LAD subset2 H3K9ac, H3K27ac, H3K4me1, and H3K4me3 are even slightly higher than in the set of all iLADs.

Functionally, both gene expression and DNA replication timing in LAD subset2 are intermediate between the set of all LADs and the set of all iLADs in all four cell lines (Fig. 5g-h, Fig. S5e-f).

### Genome-wide probing of PCH interactions with other heterochromatin regions

Heterochromatin regions have been reported to show significantly elevated association with chromocenters in mouse cells; whether similar heterochromatin interactions occur in humans which have smaller PCH remains unknown.

In addition to CENP-B TSA-seq peaks over centromeres and pericentric heterochromatin, we also observe smaller “satellite” CENP-B TSA-seq peaks that align with local peaks of nucleolar and/or lamina TSA-seq (Fig. 6a, green highlights), fall mostly within ~10-40 Mbp from the centromeres, and diminish in amplitude with increasing centromere distance. These satellite peaks are more prevalent in HCT116 and on longer chromosomes overlap with many of the LAD subset1 nucleolar TSA-seq peaks (Fig. 6a). There are a few exceptions to this trend in which these CENPB satellite peaks extend for example over the entire Chr8 p-arm or even over much of entire smaller chromosomes (Fig. 6a, Fig. 2, SFig. 2). Additionally, on the Chr 9 q-arm, immediately flanking the centromere are several narrow CENP-B peaks that align with similarly narrow MKI67IP peaks (Fig. S6b).

**Figure 6:**
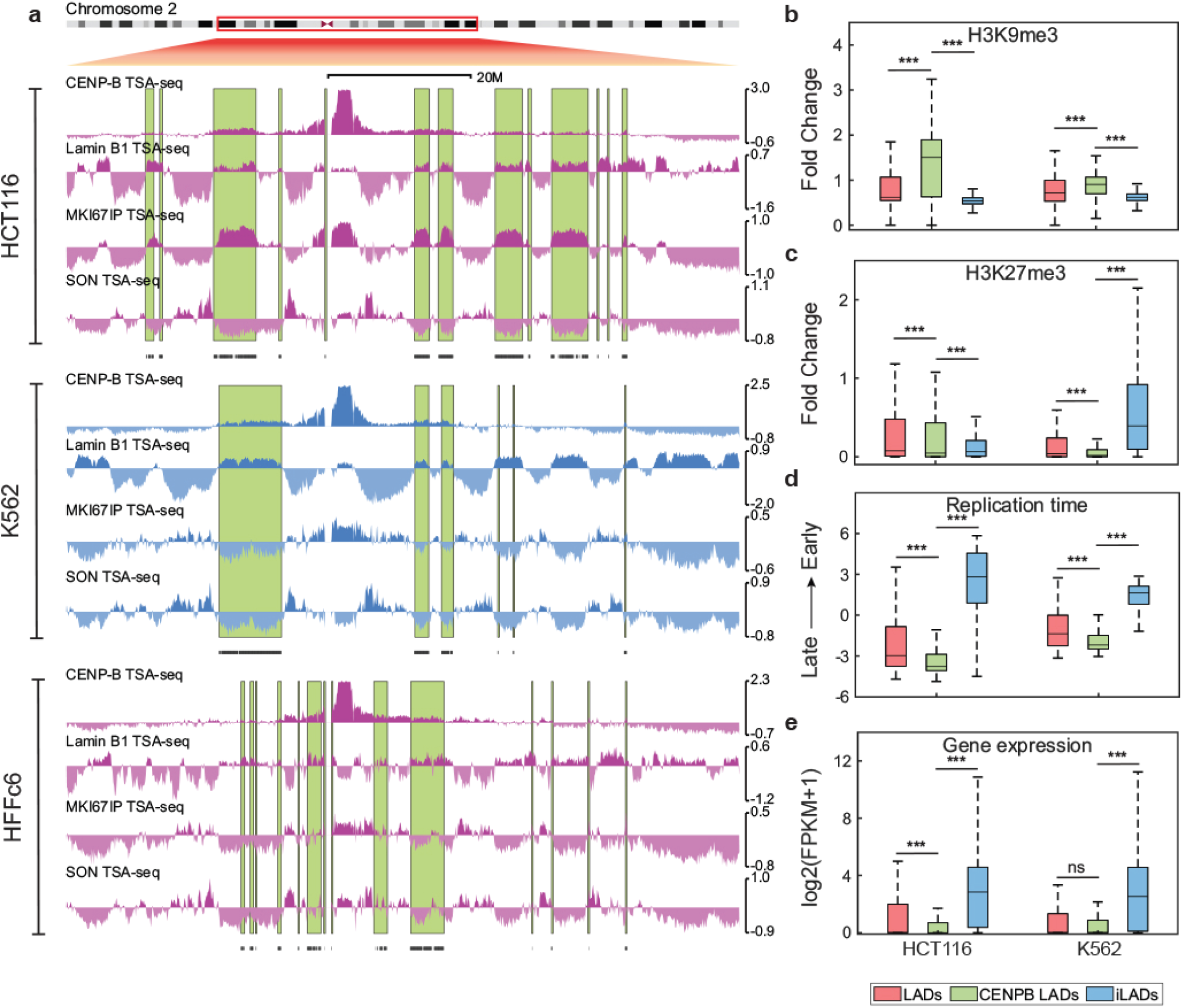
LADs with elevated PCH co-localization show later DNA replication timing and elevated H3K9me3. **a.** Top to bottom: CENP-B versus lamin B1, MKI67IP, and SON TSA-Seq and segmented bins positive in both CENP-B and lamin B1 TSA-Seq (black lines) in HCT116, K562, and HFF cells. Prominent CENP-B TSA-seq peaks over centromere regions are flanked by smaller satellite peaks (green highlights) which align with peaks in lamin B1 TSA-Seq. **b.** Segmented LADs enriched in both lamin B1 and CENP-P TSA-Seq are enriched in H3K9me3 (b), are depleted or have similar levels of H3K27me3 (c), replicate later in S-phase (d) and have similar gene expression (e) compared to all LADs in both HCT116 and K562 cells.

We defined these CENPB satellite peaks by selecting genomic regions with positive lamin B1 and low but positive CENPB TSA-seq scores (Fig. 6a, Fig. S7a-b, Table S1) (see Materials and Methods). Comparing this set of CENP-B TSA-seq satellite peaks with the set of all LADs (Fig. 6, Fig. S6), we see similar attributes to those described for LAD subset1, including elevated H3K9me3 especially in HCT116, lower H3K27me3, lower “active” histone marks, later DNA replication timing, and lower gene expression relative to the set of all LADs (Fig. 6b-e, Fig. S7b-j). The sets of CENPB satellite peaks show significant overlap across cell types and with LAD subset 1 in HCT116 (Fig. S7k-l).

The simplest interpretation is that these CENP-B TSA-seq satellite peaks are created by the colocalization of both centromeres and other heterochromatin regions at the nucleolar and/or nuclear periphery, consistent with the general alignment of these CENP-B TSA-seq peaks with MKI67IP and/or LMNB1 TSA-seq peaks. Nucleolar-associated LADs flanking the cis-linked centromere would be expected to show the highest proximity to the adjacent centromere colocalized at the nucleolar periphery, explaining the falling CENP-B TSA-seq peak amplitudes more distal on the chromosome.

### Nuclear speckle-associated, active chromosome regions map close to nucleoli in some cell lines

Given the known association of heterochromatin with the nucleolar periphery, we were surprised to see increased nucleolar TSA-seq signals over SON TSA-seq peaks that correspond to gene expression “hot-zones” ([37, 41]) (Fig. 7a). In HCT116 cells, this effect is subtle. Whereas SON TSA-seq peaks align with MKI67IP (GC) TSA-seq valleys, we frequently observe peak-within-valley local maxima for the POL1RE (FC) TSA-seq (Fig. 7a, yellow rectangles). In other cell lines, however, we see local maxima for both MKI67IP and POL1RE TSA-seq aligning with SON TSA-seq local maxima (Fig. 7a). In K562 cells, we see relatively large POL1RE TSA-seq peaks aligned with SON TSA-seq peaks.

**Figure 7:**
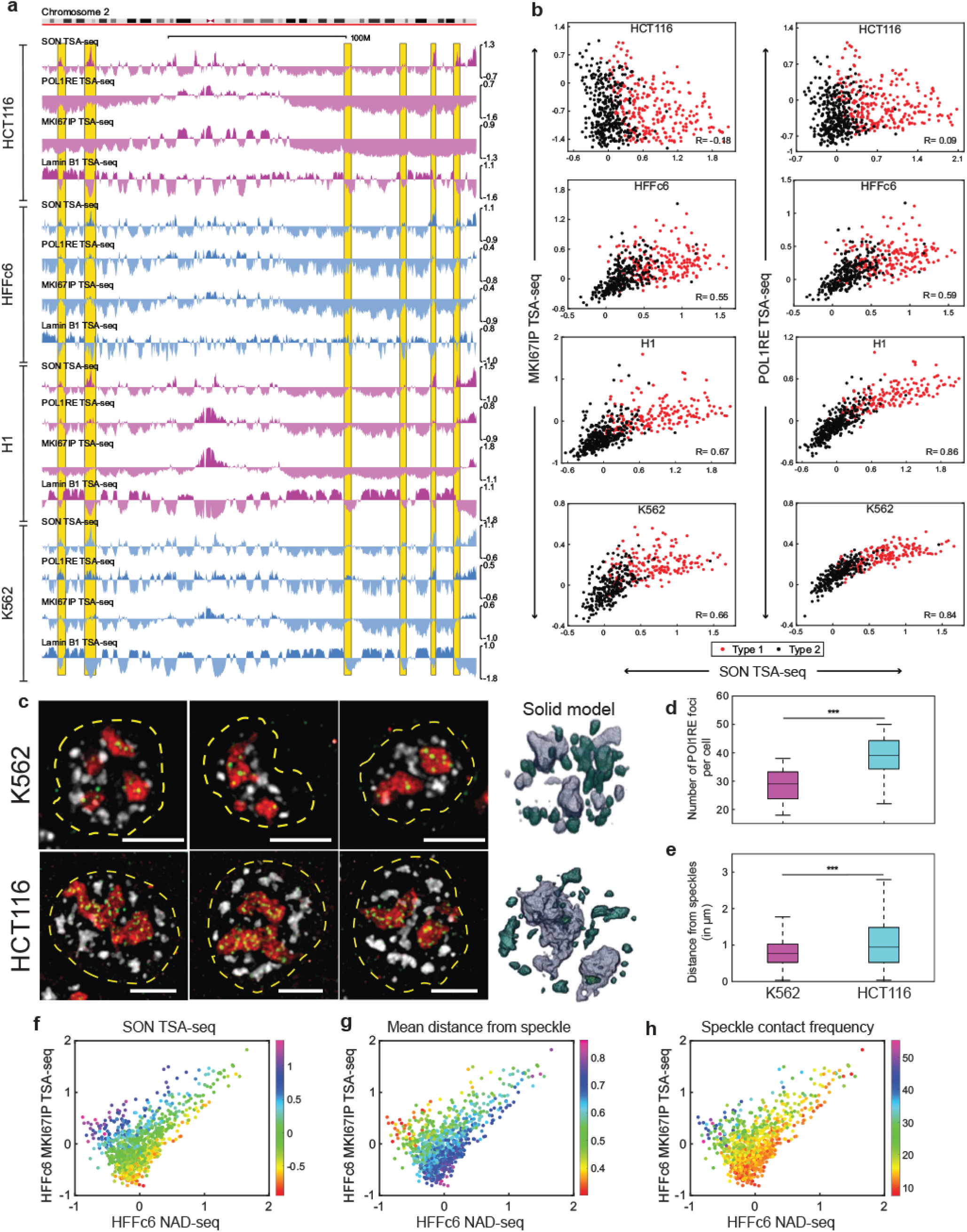
Cell-type specific relative positioning of nucleoli and nuclear speckles leads to nearby positioning of highly active chromosomal regions near nucleoli in some cell types. **a.** Pol1RE and, to a lesser extent, MKI67IP TSA-Seq peaks align with SON TSA-seq peaks (yellow highlights) especially in K562 but also H1 and HFF but not HCT116 cells. Top to bottom: SON, POL1RE, MKI67IP, and lamin B1 TSA-Seq repeated in HCT116, HFFc6, H1, and K562 cells. **b.** Scatterplots showing SON (x-axis) versus MKI67IP (left column) or POL1RE (right column) (y-axis) TSA-seq scores for Type 1 (red) versus Type 2 (black) SON TSA-seq peaks. A strong linear correlation is observed in K562 and H1, especially for SON versus POL1R1, but weaker correlation in HFF and little correlation in HCT116 cells. **c.** Immunostaining (left 3 panels) shows clustering of nuclear speckles (SON, gray) near nucleoli (MKI67IP, red; POL1R1, green; nucleus outline (from DAPI), yellow dotted lines) in K562 (top row) but not HCT116 (bottom row) cells; solid model of nucleolar (MKI67IP, grey) versus nuclear speckle (SON, green) in K562 (top) versus HCT116 (bottom) cells (right panel). Scale bar= 5um. **d-e.** Number of POL1R1 foci (y-axis) is higher in HCT116 versus K562 cells (d) but distance of nuclear speckles to nearest POL1R1 foci is significantly closer in K562 versus HCT116 cells. Measurements for box plots are from 25 nuclei for each cell type. **f-h.** Genomic regions with moderate MKI67IP TSA-Seq and low NAD-Seq scores in HFFc6 cells have high nuclear speckle association. Scatterplots of MKI67IP TSA-seq (y-axis) versus NAD-seq (x-axis) for genomic regions corresponding to FISH probes with color superimposed with SON TSA-Seq (HFFc6) (f) or mean distance from speckle (g) or contact frequency with (h) speckle (in IMR90) cells.

Plotting Type 1 (large peaks) and Type 2 (small peaks and peaks-within-valleys) SON TSA-seq local maxima scores versus nucleolar TSA-seq scores reveals a pronounced linear correlation with FC and to lesser extent GC nucleolar TSA-seq in HFF and especially K562 and H1 cells (Fig. 7b). Little correlation is observed in HCT116 cells.

Nuclear speckles noticeably clustered closer to nucleoli in K562 versus HCT116 cells (Fig. 7c-e), suggesting elevated nucleolar TSA-seq scores over SON TSA-seq peaks might be due to close positioning of nucleoli near nuclear speckles but without significant direct contact between nuclear speckles and nucleoli. Comparing the multiplexed immuno-FISH data from IMR90 fibroblasts ([45] with SON and nucleolar TSA-seq in HFF fibroblasts provides additional strong support for this explanation.

Superimposing HFF SON TSA-seq color-coded scores over HFF fibroblast NAD-seq versus MKI67IP TSA-seq scatterplots, reveals that the genomic regions with the highest SON TSA-seq scores (Fig. 7f), the lowest mean distances to the nearest nuclear speckle (Fig. 7g), and the highest nuclear speckle contact frequencies (Fig. 7h) show the lowest NAD-seq scores but have disproportionally high nucleolar TSA-seq scores. We suspect the relatively higher FC versus GC TSA-seq values over the SON TSA-seq peaks specifically in K562 cells is due to the higher proportion of FCs located near the nucleolar surface in K562 cells. Localization of FCs towards the nucleolar periphery is associated with nucleolar stress [50, 51].

Finally, in HFF cells we observe a small set of chromosomal loci at a subset of subtelomeric regions where large nuclear speckle peaks align with unusually large nucleolar TSA-seq peaks (Fig. S8).

### Interaction of centromeres with nuclear speckles in H1 cells

A second, unexpected association between nuclear speckle-associated, active chromosomal regions and a heterochromatic nuclear compartment was suggested by the alignment of smaller CENP-B TSA-seq peaks with SON TSA-seq peaks specifically in H1 cells. (Fig. 8a, yellow highlights). Scatterplots revealed a linear correlation between CENP-B and SON TSA-seq scores in H1 but not HCT116 cells (Fig. 8b-c). This correlation could reflect a lower average distance of nuclear speckles to centromeres and/or a very close contact of a fraction of centromeres with nuclear speckles.

**Figure 8:**
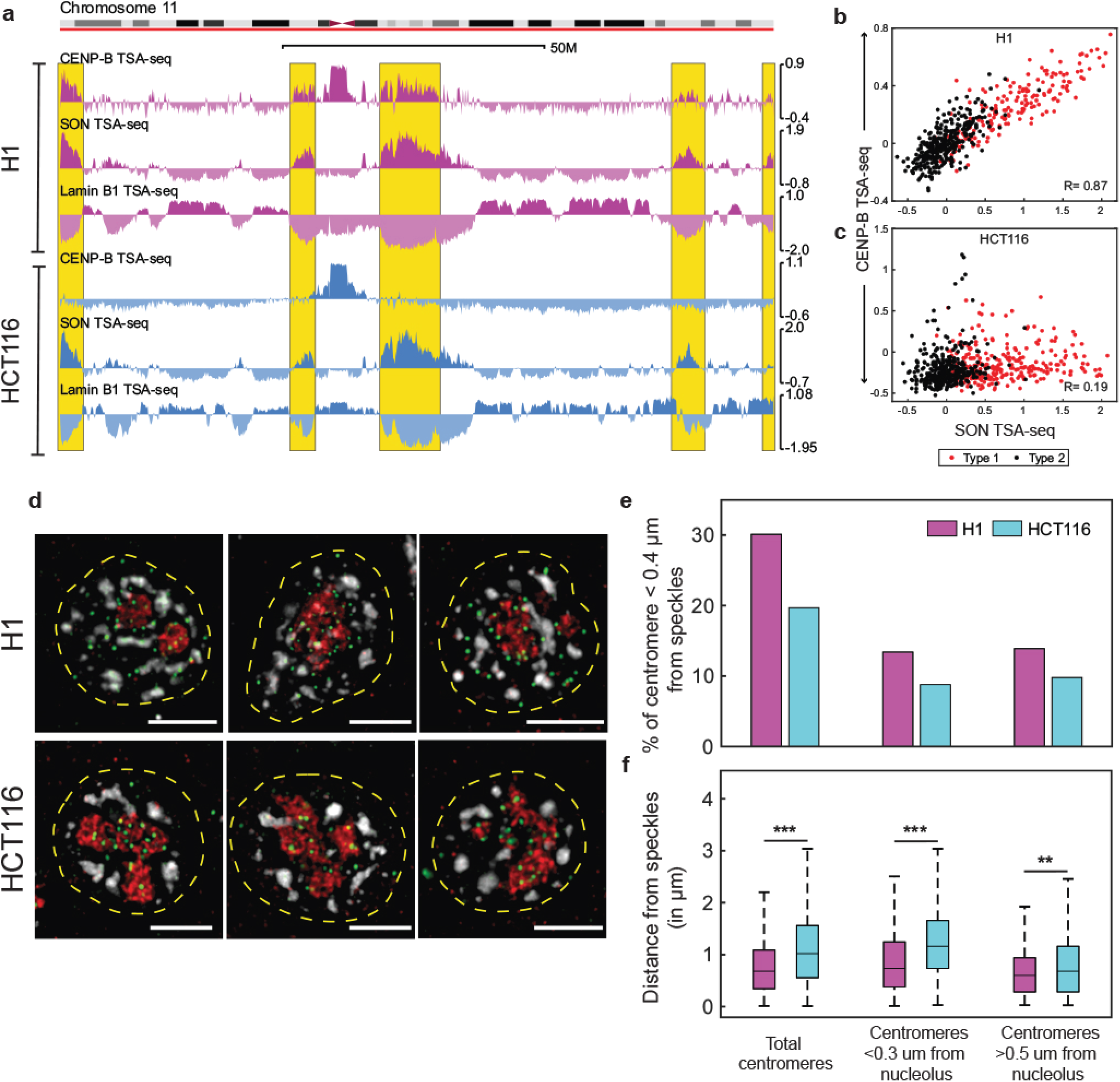
PCH and centromeres interact with nuclear speckles in H1 cells. **a.** Top to bottom: CENP-B, SON, and lamin B1 TSA-Seq repeated for H1 versus HCT116 cells: in addition to large CENP-B TSA-seq peaks over centromeres, smaller “satellite” CENP-B TSA-seq peaks are present over genomic regions (yellow highlights) aligned with SON TSA-seq peaks in H1 but not HCT116. **b-c.** Scatterplots showing SON (x-axis) versus CENP-B (y-axis) TSA-seq scores for Type 1 (red) versus Type 2 (black) SON TSA-seq peaks. A strong linear correlation is observed between SON and CENP-B TSA-Seq in H1 cells (b) but not HCT116 cells (c) for Type 1 (red) and Type 2 (black) SON TSA-seq peaks. **d-f.** Immunostaining (d) shows a larger fraction of CENP-A foci (green) in closer proximity to nuclear speckles (gray) in H1 versus HCT116 cells and this is true both for centromeres near and away from nucleoli (MKI67IP, red) (scale bars = 5um): 30% versus 20% of all centromeres are <0.4um from speckles (e) and show reduced mean distances to nearest nuclear speckle (f) in H1 versus HCT116 (n=50 cells); 13% versus 9% of centromeres <0.3um from nucleolus are <0.4um from nuclear speckles (e) and show reduced mean distances (f) to nearest nuclear speckle in H1 versus HCT116; 14% versus 10% of centromeres > 0.5um from nucleolus are < 0.4um from nuclear speckles (g) and show reduced distance (j) to nearest nuclear speckle in H1 versus HCT116 (n=25 cells for e-f).

Immunostaining shows more examples of centromeres localized both away from and near nucleoli that appear in contact with nuclear speckles in H1 versus HCT116 cells (Fig. 8d-e). There is also a significant decrease in mean distance of centromeres from the nearest nuclear speckle in H1 versus HCT116 cells (Fig. 8f). We showed above that nuclear speckles appear to cluster closer to nucleoli, but without significant contact, in K562 versus HCT116 cells (Fig. 7) and that there is significant association of centromeres with nucleoli in both cell types (Fig. 3). Thus, the CENP-B TSA-seq peaks aligned over SON TSA-seq peaks in H1 cells is likely a combined effect of both the closer proximity of nucleoli, with their associated centromeres, to nuclear speckles in H1 versus HCT116 cells plus the larger fraction of centromeres away from nucleoli specifically associating with nuclear speckles in H1 versus HCT116 cells.

## Discussion

### Summary

Here we extended TSA-seq to map the mean cytological proximity of chromosomes genome-wide relative to nucleoli and pericentric heterochromatin (PCH) in four human cell lines. Our CENPB TSA-seq represents the first mapping of genome-wide interactions with the PCH in human cells. While genomic interactions with nucleoli have been previously mapped using alternative methodologies, validation of these results has been minimal. Here we used a multiplexed immuno-FISH dataset, together with NAD-seq, to demonstrate that nucleolar TSA-seq accurately reports on mean distance of chromosome loci to nucleoli in contrast to other methods such as nucleolar DamID which do not.

Nucleolar and PCH TSA-seq reveals several novel findings. Nucleolar association increases for chromosome arms below a critical length. LADs vary in their relative association with either the nuclear lamina or nucleolus, which partially correlates with chromosome arm position but also varies across cell lines. A H3K9me3-enriched subset of LADs which replicate later and show lower gene expression than LADs as a group was identified through their increased nucleolar association specifically in HCT116 cells. A H3K27me3-enriched subset of LADs which replicate earlier and show higher gene expression than LADs as a group was identified through their association with nucleoli in K562 and H1 cells versus the nuclear lamina in HFF and HCT116 cells. Interestingly, nucleolar TSA-seq revealed significant variation across cell lines in the positioning of nuclear speckles relative to nucleoli which may explain why cell lines differ in the degree to which gene expression varies with genome radial position. PCH appears to have a minor role in the nuclear organization of heterochromatin in human cells, in contrast to the apparent situation in mouse cells. However, centromeres associate with nuclear speckles specifically in ESCs.

### Comparison with other methodologies

Building on initial comparison with several known features of nuclear genome organization, we then extensively validated the nucleolar and PCH TSA-seq through selected microscopy testing of specific TSA-seq predictions as well as through a more comprehensive genome-wide comparison with both NAD-seq and multiplexed immuno-FISH data [45].

We confirmed that nucleolar TSA-seq provides a readout of cytological-scale proximity rather than a measure of contact frequency as provided by NAD-seq. Thus, nucleolar TSA-seq and NAD-seq provide complementary measures of genome organization relative to nucleoli. Some NAD-seq datasets in the literature, however, instead show disproportionally high signals over most LADs [35]. This may be the result of co-fractionation of nuclear lamina with nucleoli in some preparations given the known attachment of nucleoli to the nuclear periphery in many cell types [52, 53]. We anticipate that nucleolar TSA-seq as compared to NAD-seq will be more easily adapted to a range of cell types, as compared to NAD-seq, as it only requires optimization of immunostaining as compared to optimization of nucleolar biochemical fractionation for each new sample. In contrast, nucleolar DamID using the 4XAP3 tethering peptide showed poor correlation with actual distance measurements of ~1000 genomic regions to the nearest nucleolus in human fibroblasts as well as both nucleolar TSA-seq and NAD-seq, correlating instead with both LMNB1 DamID and TSA-seq. The artifactually increased 4XAP3-nucleolar DamID signal over most LADs may reflect the disproportionate contribution of the small fraction of Dam methylase fusion protein localized at the nuclear lamina which is in closer molecular proximity to DNA than the larger fraction of nucleolar-localized Dam methylase.

One unique feature of nucleolar TSA-seq relative to other mapping approaches is its ability to also measure the positioning of other nuclear bodies relative to nucleoli. Comparison of nuclear speckle and nucleolar TSA-seq suggested a clustering of nuclear speckles surrounding nucleoli, whose extent varied across cell lines. This clustering was then confirmed in human fibroblasts by comparison with multiplexed immuno-FISH data [45].

### Novel biological insights

First, we showed a trend of increased nucleolar association of smaller non-NOR containing chromosomes, building on previous observations of smaller chromosomes showing a more interior localization [47]. But we showed a stronger and nonlinear correlation of nuclear association frequency with the lengths of each chromosome arm, demonstrating an increased nucleolar association of chromosome arms of non-NOR containing chromosomes below a critical arm length (~36-46 Mbp depending on cell type). Second, we confirmed how the fraction of centromeres associating with nucleoli varies significantly across cell lines, while showing how the association of centromeres with nucleoli varies among different cell lines from near constant (HCT116, H1) to quite variable (HFF, K562) frequencies among different non-NOR containing chromosomes. Third, whereas in mouse cells repressed genes and LADs have been demonstrated to have significant associations with the PCH, our CENP-B TSA-seq suggests that in human cells the PCH does not similarly act as a significant hub for heterochromatin; this likely reflects the notably smaller blocks of pericentric heterochromatin flanking human versus mouse chromosomes.

Fourth, we showed how LADs vary in their relative localization to the nuclear lamina versus nucleolar periphery. We were able to identify two subsets of LADs with distinct biochemical and functional properties based solely on changes in this differential localization across cell types. Significantly, these biochemical and functional differences between LAD subsets 1&2 were observed to varying degrees across all four cell lines independent of their nuclear localization. Thus, LAD subsets 1 and 2 may form distinct types of heterochromatin in all 4 cell lines, at least partially independent of their differential intranuclear positioning across cell types.

Fifth, we described an increased association of PCH with nuclear speckles specifically in H1 hESCs based on CENP-B TSA-seq and supported by microscopy. These results are consistent with and help explain previous 4C analysis showed the association of highly active chromosome regions with PCH specifically in mouse ESCs (mESCs) but not somatic cell lines [36]. High transcription levels of major satellite repeats and less condensed PCH, dependent on NANOG expression, have been reported in mESCs [54, 55]. We speculate that nuclear speckle association of a significant fraction of centromeres and PCH may correlate with and even possibly contribute to high PCH repeat transcription in mouse and human ESCs.

## Conclusion

Here we established nucleolar and PCH TSA-seq as new methods capable of providing new insights into nuclear genome organization. In a companion paper, we further extend this analysis of nuclear genome organization across the same four human cell lines, combining examination of nucleolar, PCH, nuclear speckle, and nuclear lamina TSA-seq with nuclear lamina DamID, and Hi-C, leading to additional novel insights. More generally, we anticipate that our new nucleolar and PCH TSA-seq will provide a valuable resource for future investigations into nuclear genome organization.

## Supplementary Figure Legends

**Figure S1: Identification of molecular markers and TSA-Seq condition for nucleolus and pericentric heterochromatin (PCH): a.** Tyramide free-radical produced after TSA staining with molecular markers from granular component (GC) of the nucleolus (middle) is expected to label larger radius of chromosome regions surrounding nucleolus compared to fibrillar center (FC) marker (bottom). **b.** POL1RE, Nucleolin, DDX18 and MKI67IP (green) staining all show less nucleoplasmic background than nucleophosmin staining (red). Both CENP-B (green) and CENP-A (red) immunostaining showed low nucleoplasmic background. **C.** TSA-Seq Condition C and Condition E produce similar results for (top to bottom) CENP-B, POL1RE and MKI67IP TSA-Seq. **d.** 2D scatterplots reveal high correlation between Condition C (x-axis) versus Condition E (y-axis) TSA staining for (top to bottom) CENP-B, POL1RE and MKI67IP TSA-Seq. TSA-Seq. TSA-seq scores were smoothed using a sliding window of 11 25kb bins.

**Figure S2: Comparison of CENP-B, POL1RE, MKI67IP, and Lamin B1 TSA-seq across all chromosomes in H1 cells**. Nucleome Browser views are based on Hg38 assembly. NORs containing chromosomes are highlighted in red.

**Figure S3: Further validation of nucleolar TSA-seq: comparison to nucleolar DamID, NAD-seq, and multiplexed immune-FISH distance measurements**. **a.** Comparisons (top to bottom) of lamin B1 TSA-Seq, lamin B1 DamID, 4xAP3 nucleolar DamID, NAD-Seq and MKI67IP TSA-Seq repeated in H1 and HFF cells shows poor correlation between nucleolar DamID and either NAD-seq or MKI67IP TSA-seq but strong correlation in H1 and still good correlation in HFF between NAD-Seq and MKI67IP TSA-Seq in H1 cells. **b-f.** Genome-wide correlations between nucleolar TSA-seq and both NAD-seq and nearest distance to nucleolus but not with nucleolar DamID, which instead correlates with lamin B1 TSA-seq: Scatterplots (100 kb bins) of distance (y-axis) of FISH probes to nucleolus (DFC, fibrillarin staining) (IMR90) versus POL1RE TSA-seq (x-axis) (HFF) (b), POL1RE TSA-seq versus NAD-seq in H1 (c) and HFF (d), 4xAp3 (nucleolar) DamID (y-axis) versus NAD-seq (x-axis) in HFF (e), and lamin B1 TSA-Seq (y-axis) versus 4xAP3 DamID (x-axis) in HFF. TSA-Seq, NAD-Seq and 4xAP3 DamID scores were smoothed using an 11-bin sliding window.

**Figure S4: Increased nucleolar association with decreasing chromosome size deconvolved into increased nucleolar association of chromosome arms below critical length-additional cell lines. a.** NOR containing chromosomes (highlighted in red, x-axis) and smaller chromosomes (Chr 16-20) have higher MKI67IP TSA-Seq scores (y axis) compared to larger chromosomes (Chr 1-12) in HFFc6 cells. **b-c.** Chromosome arm lengths less than 36 Mbp in K562 (b) and H1 (c) show higher MKI67IP TSA-Seq scores (y axis).

**Figure S5: LAD subsets identified based on their cell-type specific nuclear localization show distinctive histone marks, gene expression levels, and DNA replication timing-additional cell line and histone mark data. a-b.** Scatterplots showing MKI67IP TSA-Seq (a) and lamin B1 (b) TSA-Seq scores for LAD subset1 (yellow) and LAD subset2 (orange) using TSA-seq score selection criteria defined in Materials and Methods. **c-f.** In HFFc6 cells, LAD subset1 is enriched in H3K9me3 (c) and depleted in H3K27ac (d) relative to all LADs while LAD subset2 shows enrichment of H3K9me3 (c) and H3K27ac (d) relative to all LADs, DNA replication timing is later in LAD subset1 relative to all LADs while DNA replication timing in LAD subset2 is intermediate between all LADs and iLADs (e), and gene expression levels are lower in LAD subset1 relative to all LADs while LAD subset2 shows gene expression levels similar to all LADs but lower than all iLADs (f). **g-j.** Comparisons of histone marks H3K9ac (e), H3K27ac (f), H3K4me1 (g), and H3K4me3 (h) over LAD subset1 and subset2 relative to all LADs versus all iLADs in HCT116, H1, and K562 cells. There is an overall trend of decreased levels of these marks associated with active chromatin in LAD subset1 relative to the set of all LADs but increased levels relative to the set of all LADs for LAD subset2.

**Figure S6: Satellite CENP-B TSA-Seq peaks show varying levels of spread over chromosome arms. a-b.** In HCT116 cells, CENP-B TSA-Seq peaks (yellow highlights) overlap with nucleolar MI67IP TSA-Seq peaks over the length of the chromosome 8 p-arm (top) but only over the chromosome 9 p-arm region flanking the centromere, despite the presence of multiple MKI67IP TSA-seq peaks over most of the Chr 9 p-arm length (bottom).

**Figure S7: PCH-interacting LADs show distinctive biochemical and functional properties a.** Scatterplots showing lamin B1 and CENP-B TSA-Seq scores for LAD regions with increased interactions with pericentric heterochromatin (PCH) but lying outside of centromere regions (“CENP-B LADs”, green) in HCT116 (left), K562 (middle), and HFF (right) cells (see Materials and Methods for TSA-seq score selection criteria). **b-f.** CENP-B LADs in HFF cells show slightly elevated H3K9me3 (b), lower H3K27ac (c), similar H3K4me3 (d), slightly later DNA replication (e), and lower gene expression (f) relative to the set of all LADs; levels over the set of all iLADs are shown for comparison. **g-j** CENP-B LADs in HCT116 and K562 cells show lower H3K9ac (g), lower H3K27ac (h), lower H3K4me1 (i), and lower (HCT116) or similar (K562) H3K4me3 (j) compared to the set of all LADs; levels over the set of all iLADs are shown for comparison. **k.** Comparison of CENP-B LADs in HCT116, K562, and HFFc6 cells using Venn-diagrams; units are numbers of 25 kb genomic bins segmented in (a) in common among different cell line comparisons. **l.** Overlap in HCT116 cells between LAD subset1, CENP-B LADs, and set of all LADs shown using Venn-diagram; units are numbers of 25 kb genomic bins overlapping among these categories. Note that set of all LADs was defined by DamID whereas the set of CENP-B LADs was defined as in (a).

**Figure S8: In HFFc6 cells a subset of subtelomeric regions have both high nucleolus and SON TSA-Seq score. a-g.** Genome browser views comparing (top to bottom) lamin B1 DamID, lamin B1 TSA-Seq, lamin B1 DamID, MKI67IP TSA-Seq, POL1RE TSA-Seq and SON TSA-Seq peaks. A small subset of subtelomeric regions (yellow highlights) have the unusual combination of showing large peaks in both nucleolus and SON TSA-Seq and mostly valleys in lamin B1 DamID and TSA-seq.

## Methods

### Cell culture

H1-ESC (WA01), HFF-hTert-Clone 6 cells and HCT116 were obtained thorough the 4D Nucleome Consortium and were cultured according to the 4DN SOPs (https://www.4dnucleome.org/cell-lines.html). K562 cells were obtained from the ATCC and were cultured according to the ENCODE Consortium protocol (http://genome.ucsc.edu/ENCODE/protocols/cell/human/K562_protocol.pdf).

### Antibodies

The following primary antibodies were used in this study: anti-MKi67IP (1:2000, Atlas Antibodies, catalog no. HPA035735), anti-POL1RE (1:2000, Atlas Antibodies, catalog no. HPA052400), anti-DDX18 (1:2000, Atlas Antibodies, catalog no. HPA041056), anti-Nucleolin (1:1000, Abcam, catalog no. ab70493), anti-Nucleophosmin (1:1000, Abcam, catalog no. ab86712), anti-CENP-B (1:800, Santa Cruz Biotechnology, Inc, catalog no. sc-22788), anti-CENP-A (1:1000, Abcam, catalog no. ab13939), custom-made anti-SON (Chen et.al., 2018; 1:2000; Pacific Immunology, catalog no. PACIFIC10700), and anti-LMNB1 (1:700, Abcam, catalog no. ab16048). All primary antibodies were rabbit polyclonal except for the anti-Nucleoplasmin and anti-CENP-A which were mouse monoclonal.

The following secondary antibodies were used in this study: goat anti-rabbit HRP (1:1000, Jackson ImmunoResearch, catalog no. 111-035-144), streptavidin-Alexa Fluor 594 (1:200, Invitrogen, catalog no. S-11227), goat anti-rabbit FITC (1:250 or 1:500, Jackson ImmunoResearch, catalog no. 111-095-144), goat anti-mouse Alexa Fluor 594 (1:250 or 1:500, Jackson ImmunoResearch, catalog no. 115-585-146), streptavidin-HRP (1:10,000, Invitrogen, catalog no. 43-4323).

### Immunostaining

Cells were plated on coverslips and were harvested at ~80% confluency. Cells were rinsed with PBS and then fixed in PBS for 20 min at room temperature (RT) with 1.6% paraformaldehyde (P6148, Sigma-Aldrich). Cells were then rinsed with PBS and then permeabilized with 0.5% Triton X-100 (T8787, Sigma-Aldrich) in PBS at RT for 3 x 5 min. Cells were then rinsed with rinsed with 0.1% Triton X-100 in PBS (0.1%PBST) and then blocked with blocking buffer consisting of 5% normal goat serum (G9023, Sigma-Aldrich) in 0.1% PBST for 1hr at RT. Cells were then incubated with appropriate dilutions of primary antibodies in blocking buffer at 4°C for 12 hrs. Cells were then washed with 0.1% PBST at RT for 3 x 5 mins and incubated with appropriate dilutions of secondary antibodies in blocking buffer at 4°C for 10 hrs. Cells were washed with 0.1% PBST at RT for 3 x 5 mins. Coverslips were mounted in DAPI containing, anti-fading medium (0.3 μg/ml DAPI [Sigma-Aldrich]/10% w/v Mowiol 4-88[EMD Millipore]/1% w/v DABCO [Sigma-Aldrich]/25% glycerol/0.1 M Tris, pH 8.5).

### Primary antibody labeling and immunostaining

For doing triple immunostaining, primary antibodies were first labelled with fluorescent dyes using the Mix-n-Stain antibody labelling kit (Biotium) according to the manufacturer’s protocol. anti-POL1RE was linked to CF488 (92273, Biotium), anti-MKi67IP was linked to CF594 (92276, Biotium) and anti-SON was linked to CF640R (92278, Biotium). These labelled antibodies were used for immunostaining as described above with the following modification. The cells were then incubated with labelled primary antibodies in blocking buffer at 4°C for 12 hrs. After incubation, cells are then washed with 0.1% PBST at RT for 3 x 5 mins. Coverslips were mounted in DAPI containing, anti-fading medium (0.3 μg/ml DAPI [Sigma-Aldrich]/10% w/v Mowiol 4-88[EMD Millipore]/1% w/v DABCO [Sigma-Aldrich]/25% glycerol/0.1 M Tris, pH 8.5).

### TSA-Seq

Nucleolar and PCH TSA-Seq was performed using either Condition C (labeling with 1:3000 tyramide biotin, 50% sucrose and 0.0015% hydrogen peroxide) or Condition E (labeling with 1:300 tyramide biotin, 50% sucrose and 0.0015% hydrogen peroxide) [41] with the following minor modification: 150ul of Dynabeads M-270 streptavidin (Invitrogen, catalog no. 65306) was used to purify the biotinylated DNA. LMNB1 TSA-seq was performed using Condition AI for HFFc6, H1, and K562 and Condition A for HCT116 cells [41].

### NAD-seq

#### Nucleoli isolation for NAD-seq

H1 cells were grown in ten 10 cm plates until they reached 80-90% confluence, for a total of 30-100 x 10^6^ cells per preparation. HFF-hTERT clone 6 cells (passages 22 and 23) were grown to 80-90% confluency in thirteen 150 mm tissue culture dishes. Old media was removed, and fresh media was added 1 h prior to nucleoli preparation. An additional plate grown in parallel was reserved for total genomic DNA extraction (Quick-DNA Universal Kit (Zymo Research, CA)). Formaldehyde-crosslinked cells were prepared as described ([34, 46], adapted from [32, 56]). Briefly, cells were fixed by the addition of 1% formaldehyde added directly to the media and incubated at room temperature for 10 minutes. Media was removed, and cross-linking was quenched by adding 5 ml 1M glycine. The cells were washed with ice-cold PBS, collected by scraping in 40 ml PBS, and centrifuged at 200 x g for 5 min at 4°C. The cell pellet was re-suspended in 1 ml high magnesium (HM) buffer (10 mM HEPES-KCl pH 7.5, 0.88 M sucrose, 12 mM MgCl2, 1 mM DTT). Cells were then sonicated on ice (12-16 bursts for 10s at full power for HFF and 15 bursts for H1 cells) using a Soniprep 150 (MSE) with a fine probe. The release of nucleoli was monitored microscopically. Nucleoli were pelleted by centrifugation in a microfuge at 15,000 x g for 20 seconds and re-suspended in 0.5 ml low magnesium (LM) buffer (10 mM HEPES-KCl pH 7.5, 0.88 M sucrose, 1 mM MgCl2, 1 mM DTT, proteinase cocktail HALT (Thermo Scientific, #78438)). The sample was subjected to the second round of sonication (1 burst for HFF for 10s at full power; 2-3 bursts for H1 cells) and centrifuged at 15,000 x g for 20 seconds to pellet nucleoli. The pellet was resuspended in 20/2-TE buffer (20 mM Tris-Cl pH 7.5, 2 mM EDTA) for immediate use or in 20/2-TE + 50% glycerol to snap-freeze in liquid nitrogen and store at −80°C.

#### Immunoblot analyses of NADseq preparations

Primary antibodies were fibrillarin (ab5821, Abcam, Cambridge, MA) and actin (beta-actin, Sigma A1978). Horseradish peroxidase (HRP)-coupled anti-mouse and anti-rabbit secondary antibodies (Jackson ImmunoResearch, West Grove, PA) were used. Protein concentrations from total cell lysates or isolated nucleoli were assessed by Bradford assay (BioRad reagent Blue R-250, cat. #161-0436 with BSA as a standard). 10 μg of each sample were loaded per lane on 12% SDS-PAGE gels and transferred to PVDF membranes for 2 hours at 80 V, 4°C. Membranes were blocked in PBS-nonfat 5% milk and incubated with antibodies according to manufacturer instructions.

#### Quantitative PCR

DNA extraction from whole input cells and purified nucleoli was performed using a Quick-DNA Universal Kit (Zymo Research, CA). DNA concentration was measured using a Qubit dsDNA HS Assay kit (Invitrogen, Eugene, OR). 5 ng of DNA from each sample was analyzed using the Kapa Cyber Fast Q-PCR Kit (Kapa Biosystems, Wilmington, MA). The following rDNA primers (designed using Primer3Plus software) were used: Fw: gaa ctt tga agg ccg aag tg; Rv: atc tga acc cga ctc cct tt. The PCR program used was as follows: hold at 98°C for 30 s followed by 40 cycles of 95°C for 10 s and 60°C for 30 s. All signals from nucleolar samples were normalized to the signals from the input cell samples, using the 2−ΔΔ*CT* method for quantification (Life Technologies).

#### DNA isolation, deep sequencing

Total genomic and nucleolar DNA was isolated with Quick-DNA Universal Kit (Zymo Research, CA, USA). Libraries were constructed using Illuminàs TruSeq DNA PCR-free Library Preparation kit (350 bp), and fragments were size-selected by sample-purifying beads. 150bp paired-end sequencing was performed using Illumina HiSeq X Ten sequencing system. For more details on sequencing, please see the metadata files associated with the data at data.4dnucleome.org, (https://data.4dnucleome.org/belmont_lab_nucleolus_centromere_TSA-seq)

### TSA-seq and NAD-seq data processing

We used the pipeline as described in previous paper (Zhang et.al 2021) to process the TSA-seq and NAD-seq data. For NAD-seq only one read of a pair was used for mapping the reads. For normalization of NAD-seq data, total genomic DNA sample and nucleolar DNA sample was used as input and pulldown respectively. For Fig 3a, POL1RE and MKI67IP TSA-Seq, raw sequencing reads were mapped to the Telomere-to-Telomere (version 1.1).

### Genome segmentation to define Subset1, Subset2 and CENP-B LADs

Initial exploration of the TSA-seq data used a scatterplot tool that allowed users to select genomic bins falling simultaneously into Regions of Interest (ROI) in multiple scatterplots (https://scatterplot.nucleome.org/). For analysis purposes, we then defined specific windows of TSA-seq values to select different subsets of genomic bins. Subset1 LAD genomic bins were defined as all genomic bins with MKI67IP TSA-Seq scores <0 in H1, K562 and HFFc6 cells but >0.5 in HCT116 cells (Fig. S5a). Subset2 LADs were defined as all genomic bins that met the following two conditions simultaneously (logical AND): 1) all genomic bins with <0 Lamin B1 TSA-Seq scores in H1 and K562 cells but >0 in HCT116 and HFFc6 cells; 2) all genomic bins identified by (1) but also with >0 MKI67IP TSA-Seq scores in H1 and K562 cells but <0 in HCT116 and HFFc6 cells (Fig. S5a&b). CENP-B LADs were defined also by applying two conditions simultaneously: 1) all genomic bins with scores >0.1 for both CENP-B and lamin B1 TSA-Seq; 2) all genomic bins identified by (1) but also with <0.5 CENP-B TSA-seq scores (effectively excludes centromere regions).

### 4xAP3 Nucleolar DamID

4xAP3 DamID data was generated as previously described [30].

### Microscopy and image processing

Images of TSA staining (Fig. 1a, Fig. S1b) were collected on Personal Delta Vision microscope (GE Healthcare) equipped with a Coolsnap HQ camera and Plan Apo N 60x/1.42-NA oil-immersion objective (Olympus). 3D optical section were collected at 0.2μm z-increment. Images were deconvolved using an enhanced-ratio iterative constrained algorithm. Co-immunostaining images of nuclear bodies (Fig. 3c, Fig. 7c, Fig. 8d) were collected on OMX-V4 microscope (GE Healthcare) equipped with a U Plan S-Apo 100x/1.40-NA oil-immersion objective (Olympus) and two Evolve EMCCD camera (Photometrics). Z-sections in the 3d optical section were 0.2μm apart. The images were deconvolved using an enhanced-ratio iterative constrained algorithm.

### Image analysis

Image analysis was done using FIJI software (Image J). Fig. 3c, Fig. 7c and Fig. 8d were scaled using a gamma value of 0.5, applied uniformly across the entire image. For K562 cells (in Fig. 3c and Fig. 7c) and H1 (Fig. 3c and Fig. 8d) 2-3 optical z-sections were projected in the x-y plane using a maximum-intensity projection algorithm. To measure the distance of centromeres to either a nucleolus (Fig. 3d) or speckle (Fig 8 e-f) or POL1RE foci to speckles (Fig 7 d-e), we used an ImageJ plugin developed in the Belmont laboratory (https://github.com/omidalam/compartment_dist). Briefly, a sum projection over 5 optical z-sections is created for both the locus and the compartment channels centered around the brightest pixel of the locus. To define the locus and compartment boundary, the background (mode in pixel intensity histogram) was subtracted from the image and the boundary was defined using Full-Width at Half Maximum (FWHM) thresholding of the compartment intensities centered in a local window over the locus location-essentially segmenting the compartment using an intensity threshold 50% of the local maximum. The shortest line connecting the boundary of compartment and loci center was measured as their distance. Distances of centromeres to the nearest nuclear speckle were measured using the previously mentioned ImageJ plugin; distances of centromeres to the nearest nucleolus were measured by drawing a line from the center of the centromere to the edge of the nucleolus.

### Normalization of MKI67IP TSA-seq score over centromeres

Average MKI67IP TSA-seq scores over the ribosomal gene cluster were obtained from the tracks mapped to the T-to-T genome assembly. This score was then used to normalize the MKI67IP TSA-seq score in hg38 genome assembly. The MKI67IP TSA-seq score over the centromeres were then plotted in Fig 3b. The centromere coordinates in the hg38 genome assembly were obtained from NCBI (https://www.ncbi.nlm.nih.gov/grc/human).

### ChIP-seq data processing

Histone ChIP-seq data for four cell lines were downloaded from the ENCDOE website or 4DN Data Portal (Table S2). The genome was divided into non-overlapping 25 kb intervals, and the average fold change signal over control was calculated for each track. Next, we removed bins that completely overlapped with the blacklist region generated by ENCODE (accession ID: ENCSR636HFF) or the gap regions of hg38 provided by the UCSC Genome Browser, which represent unfinished sequences in the hg38 genome assembly.

The remaining bins were used to remove background signals as described previously [37]. To do this, we created a histogram of average fold change signals across the available bins. The mode of the histogram was considered the background signal for all histone targets except for H3K9me3. For H3K9me3, the 0.5% percentile of the signal was considered the background. The adjusted signal was then calculated by subtracting the background signal from the original signal per bin. If the adjusted signal was below zero, it was replaced with zero.

### RNA-seq data processing

Processed RNA-seq data was obtained from [41].

## Data Availability

Both primary sequencing data plus processed data, including segmentation of different genomic regions is available at: https://data.4dnucleome.org/belmont_lab_nucleolus_centromere_TSA-seq

## Author Contributions

PK designed and performed all of the nucleolar and PCH TSA-seq and microscopy experiments, collected the data, and conducted data analysis and microscopy image analysis with ASB’s guidance. PK and LZ generated the lamin B1 TSA-seq data. OG developed the distance measurement software for image analysis and assisted in data analysis. YZ processed ChIP-seq and the two fraction Repli-seq data. TvS, DPH and BvS provided the 4xAP3 DamID data. TS and DMG provided the two fraction Repli-seq data. AV and PDK provided the NAD-seq data. PK and ASB wrote the manuscript with editing contributions from PK, BvS, JM, LZ, DMG, YZ, AV, and OG. ASB supervised the overall study.

## Acknowledgements

This work was supported by the National Institutes of Health Common Fund 4D Nucleome Program grants U54DK107965 (A.S.B., B.v.S., D.M.G., and J.M.), UM1HG011593 (J.M., A.S.B., and D.M.G.), and U01CA260669 (P.D.K.). Research at the Netherlands Cancer Institute is supported by an institutional grant of the Dutch Cancer Society and of the Dutch Ministry of Health, Welfare and Sports. The Oncode Institute is partially funded by the Dutch Cancer Society. We thank Krisna Mohan Pars in the Rene Maehr laboratory (UMass) for providing H1 cells and Liyan Yang in the Job Dekker laboratory (UMass) for providing HFFc6 cells used for the NAD-seq.

## Notes

https://data.4dnucleome.org/belmont_lab_nucleolus_centromere_TSA-seq

### Competing Interest Statement

The authors have declared no competing interest.

## References

1. Brown, S.W., Heterochromatin. Science, 1966. 151(3709): p. 417–25.

2. Heitz, E., Das Heterochromatin der Moose. I. Jahrb Wiss Bot, 1928. 69: p. 762–818.

3. Passarge, E., Emil Heitz and the concept of heterochromatin: longitudinal chromosome differentiation was recognized fifty years ago. Am J Hum Genet, 1979. 31(2): p. 106–15.

4. Bank, E.M. and Y. Gruenbaum, The nuclear lamina and heterochromatin: a complex relationship. Biochem Soc Trans, 2011. 39(6): p. 1705–9.

5. van Steensel, B. and A.S. Belmont, Lamina-Associated Domains: Links with Chromosome Architecture, Heterochromatin, and Gene Repression. Cell, 2017. 169(5): p. 780–791.

6. Belmont, A.S., Nuclear Compartments: An Incomplete Primer to Nuclear Compartments, Bodies, and Genome Organization Relative to Nuclear Architecture. Cold Spring Harb Perspect Biol, 2021.

7. Padeken, J. and P. Heun, Nucleolus and nuclear periphery: velcro for heterochromatin. Current opinion in cell biology, 2014. 28: p. 54–60.

8. Bizhanova, A. and P.D. Kaufman, Close to the edge: Heterochromatin at the nucleolar and nuclear peripheries. Biochim Biophys Acta Gene Regul Mech, 2021. 1864(1): p. 194666.

9. Carvalho, C., et al., Chromosomal G-dark Bands Determine the Spatial Organization of Centromeric Heterochromatin in the Nucleus. Mol. Biol. Cell, 2001. 12(11): p. 3563–3572.

10. Vourc’h, C., et al., Cell cycle-dependent distribution of telomeres, centromeres, and chromosome-specific subsatellite domains in the interphase nucleus of mouse lymphocytes. Exp. Cell Res., 1993. 205: p. 142–151.

11. Chaly, N. and S.B. Munro, Centromeres Reposition to the Nuclear Periphery during L6E9 Myogenesisin Vitro. Experimental Cell Research, 1996. 223(2): p. 274–278.

12. Solovei, I., et al., Differences in centromere positioning of cycling and postmitotic human cell types. Chromosoma, 2004. 112(8): p. 410–23.

13. Takizawa, T., K.J. Meaburn, and T. Misteli, The meaning of gene positioning. Cell, 2008. 135(1): p. 9–13.

14. Bickmore, W.A., The spatial organization of the human genome. Annu Rev Genomics Hum Genet, 2013. 14: p. 67–84.

15. Finlan, L.E., et al., Recruitment to the nuclear periphery can alter expression of genes in human cells. PLoS Genet, 2008. 4(3): p. e1000039.

16. Zink, D., et al., Transcription-dependent spatial arrangements of CFTR and adjacent genes in human cell nuclei. J Cell Biol, 2004. 166(6): p. 815–25.

17. Csink, A.K. and S. Henikoff, Genetic modification of heterochromatin association and nuclear organization in Drosophila. Nature, 1996. 381: p. 529–531.

18. Dernburg, A.F., et al., Perturbation of nuclear architecture by long-distance chromosome interactions. Cell, 1996. 85: p. 745–759.

19. Brown, K.E., et al., Dynamic repositioning of genes in the nucleus of lymphocytes preparing for cell division. Molecular cell, 1999. 3(2): p. 207–17.

20. Brown, K.E., et al., Expression of alpha- and beta-globin genes occurs within different nuclear domains in haemopoietic cells. Nat Cell Biol, 2001. 3(6): p. 602–6.

21. Fisher, A.G. and M. Merkenschlager, Gene silencing, cell fate and nuclear organisation. Curr Opin Genet Dev, 2002. 12(2): p. 193–7.

22. Ragoczy, T., et al., Functional redundancy in the nuclear compartmentalization of the late-replicating genome. Nucleus, 2014. 5(6): p. 626–35.

23. Politz, J.C.R., D. Scalzo, and M. Groudine, The redundancy of the mammalian heterochromatic compartment. Curr Opin Genet Dev, 2016. 37: p. 1–8.

24. Politz, J.C., D. Scalzo, and M. Groudine, Something silent this way forms: the functional organization of the repressive nuclear compartment. Annual review of cell and developmental biology, 2013. 29: p. 241–70.

25. Guelen, L., et al., Domain organization of human chromosomes revealed by mapping of nuclear lamina interactions. Nature, 2008. 453(7197): p. 948–51.

26. Lund, E.G., et al., Distinct features of lamin A-interacting chromatin domains mapped by ChIP-sequencing from sonicated or micrococcal nuclease-digested chromatin. Nucleus, 2015. 6(1): p. 30–9.

27. Briand, N. and P. Collas, Lamina-associated domains: peripheral matters and internal affairs. Genome Biol, 2020. 21(1): p. 85.

28. van Schaik, T., et al., Cell cycle dynamics of lamina-associated DNA. EMBO Rep, 2020. 21(11): p. e50636.

29. Kind, J., et al., Genome-wide maps of nuclear lamina interactions in single human cells. Cell, 2015. 163(1): p. 134–47.

30. Wang, Y., et al., SPIN reveals genome-wide landscape of nuclear compartmentalization. Genome Biol, 2021. 22(1): p. 36.

31. Bersaglieri, C., et al., Genome-wide maps of nucleolus interactions reveal distinct layers of repressive chromatin domains. Nat Commun, 2022. 13(1): p. 1483.

32. Nemeth, A., et al., Initial genomics of the human nucleolus. PLoS Genet, 2010. 6(3): p. e1000889.

33. van Koningsbruggen, S., et al., High-resolution whole-genome sequencing reveals that specific chromatin domains from most human chromosomes associate with nucleoli. Mol Biol Cell, 2010. 21(21): p. 3735–48.

34. Vertii, A., et al., Two contrasting classes of nucleolus-associated domains in mouse fibroblast heterochromatin. Genome Res, 2019. 29(8): p. 1235–1249.

35. Dillinger, S., T. Straub, and A. Nemeth, Nucleolus association of chromosomal domains is largely maintained in cellular senescence despite massive nuclear reorganisation. PLoS One, 2017. 12(6): p. e0178821.

36. Wijchers, P.J., et al., Characterization and dynamics of pericentromere-associated domains in mice. Genome Res, 2015. 25(7): p. 958–69.

37. Chen, Y., et al., Mapping 3D genome organization relative to nuclear compartments using TSA-Seq as a cytological ruler. J Cell Biol, 2018. 217(11): p. 4025–4048.

38. Pluta, A.F., et al., Identification of a subdomain of CENP-B that is necessary and sufficient for localization to the human centromere. J Cell Biol, 1992. 116(5): p. 1081–93.

39. Thul, P.J., et al., A subcellular map of the human proteome. Science, 2017. 356(6340).

40. Biggiogera, M., et al., Nucleolar distribution of proteins B23 and nucleolin in mouse preimplantation embryos as visualized by immunoelectron microscopy. Development, 1990. 110(4): p. 1263–70.

41. Zhang, L., et al., TSA-seq reveals a largely conserved genome organization relative to nuclear speckles with small position changes tightly correlated with gene expression changes. Genome Res, 2020.

42. Nurk, S., et al., The complete sequence of a human genome. Science, 2022. 376(6588): p. 44–53.

43. van Sluis, M., et al., NORs on human acrocentric chromosome p-arms are active by default and can associate with nucleoli independently of rDNA. Proc Natl Acad Sci U S A, 2020. 117(19): p. 10368–10377.

44. Rodrigues, A., et al., Nucleoli and the nucleoli-centromere association are dynamic during normal development and in cancer. Mol Biol Cell, 2023. 34(4): p. br5.

45. Su, J.H., et al., Genome-Scale Imaging of the 3D Organization and Transcriptional Activity of Chromatin. Cell, 2020. 182(6): p. 1641–1659 e26.

46. Bizhanova, A., et al., Distinct features of nucleolus-associated domains in mouse embryonic stem cells. Chromosoma, 2020. 129(2): p. 121–139.

47. Bolzer, A., et al., Three-dimensional maps of all chromosomes in human male fibroblast nuclei and prometaphase rosettes. PLoS biology, 2005. 3(5): p. e157.

48. Croft, J.A., et al., Differences in the localization and morphology of chromosomes in the human nucleus. J Cell Biol, 1999. 145(6): p. 1119–31.

49. Bian, Q., et al., *beta-Globin cis-elements determine differential nuclear targeting through epigenetic modifications*. The Journal of cell biology, 2013. 203(5): p. 767–83.

50. Frottin, F., et al., The nucleolus functions as a phase-separated protein quality control compartment. Science, 2019. 365(6451): p. 342–347.

51. Latonen, L., Phase-to-Phase With Nucleoli - Stress Responses, Protein Aggregation and Novel Roles of RNA. Front Cell Neurosci, 2019. 13: p. 151.

52. Bourgeois, C.A., D. Hemon, and M. Bouteille, Structural relationship between the nucleolus and the nuclear envelope. J Ultrastruct Res, 1979. 68(3): p. 328–40.

53. Bourgeois, C.A. and J. Hubert, Spatial relationship between the nucleolus and the nuclear envelope: structural aspects and functional significance. Int Rev Cytol, 1988. 111: p. 1–52.

54. Novo, C.L., et al., The pluripotency factor Nanog regulates pericentromeric heterochromatin organization in mouse embryonic stem cells. Genes Dev, 2016. 30(9): p. 1101–15.

55. Efroni, S., et al., Global transcription in pluripotent embryonic stem cells. Cell Stem Cell, 2008. 2(5): p. 437–47.

56. Sullivan, G.J., et al., Human acrocentric chromosomes with transcriptionally silent nucleolar organizer regions associate with nucleoli. EMBO J, 2001. 20(11): p. 2867–74.

