## Supplementary figures and images for "Nucleolus and centromere TSA-Seq reveals variable localization of heterochromatin in different cell types"

### Supplementary Figures

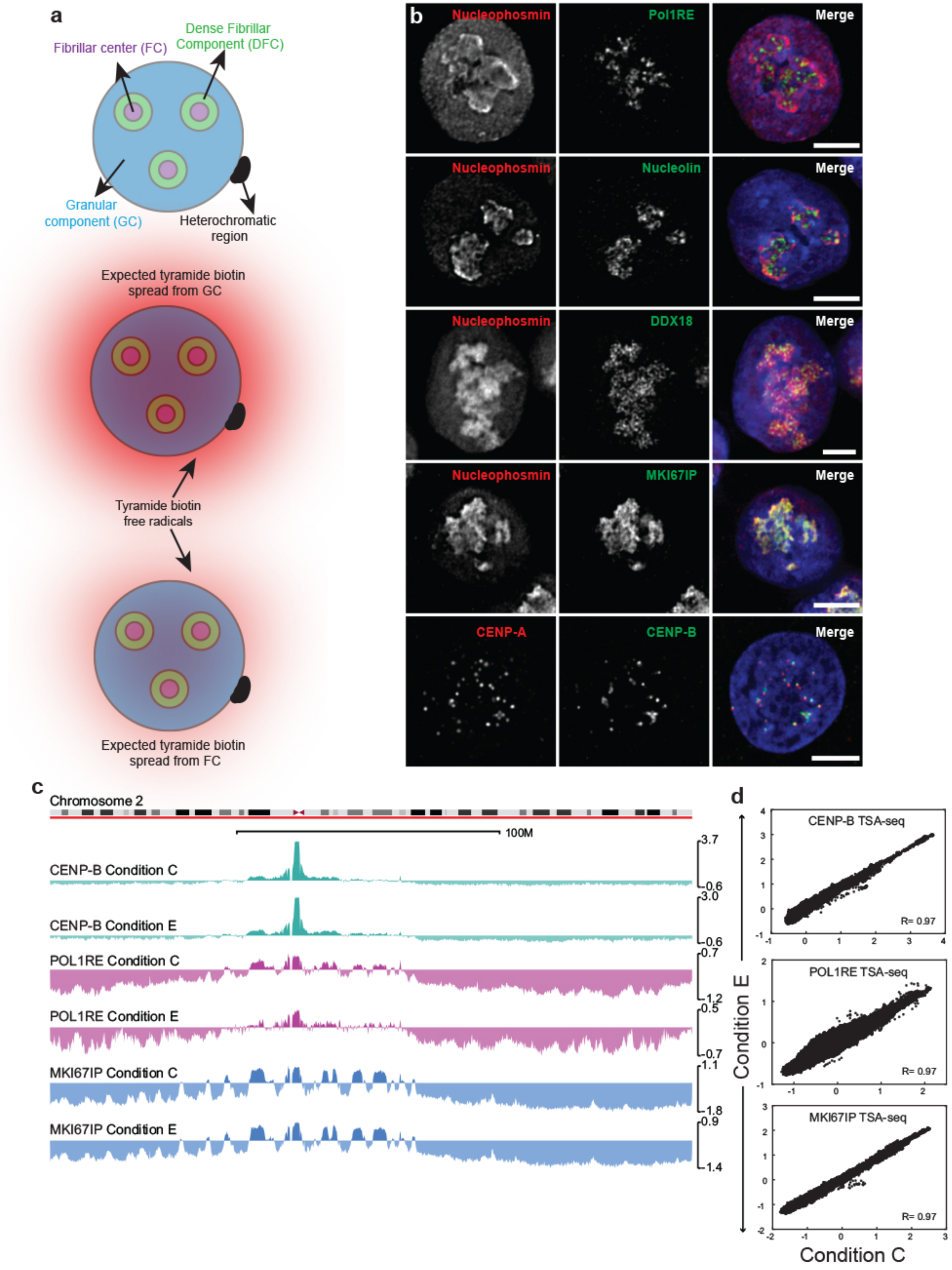

#### Figure S2

##### H1 TSA-seq

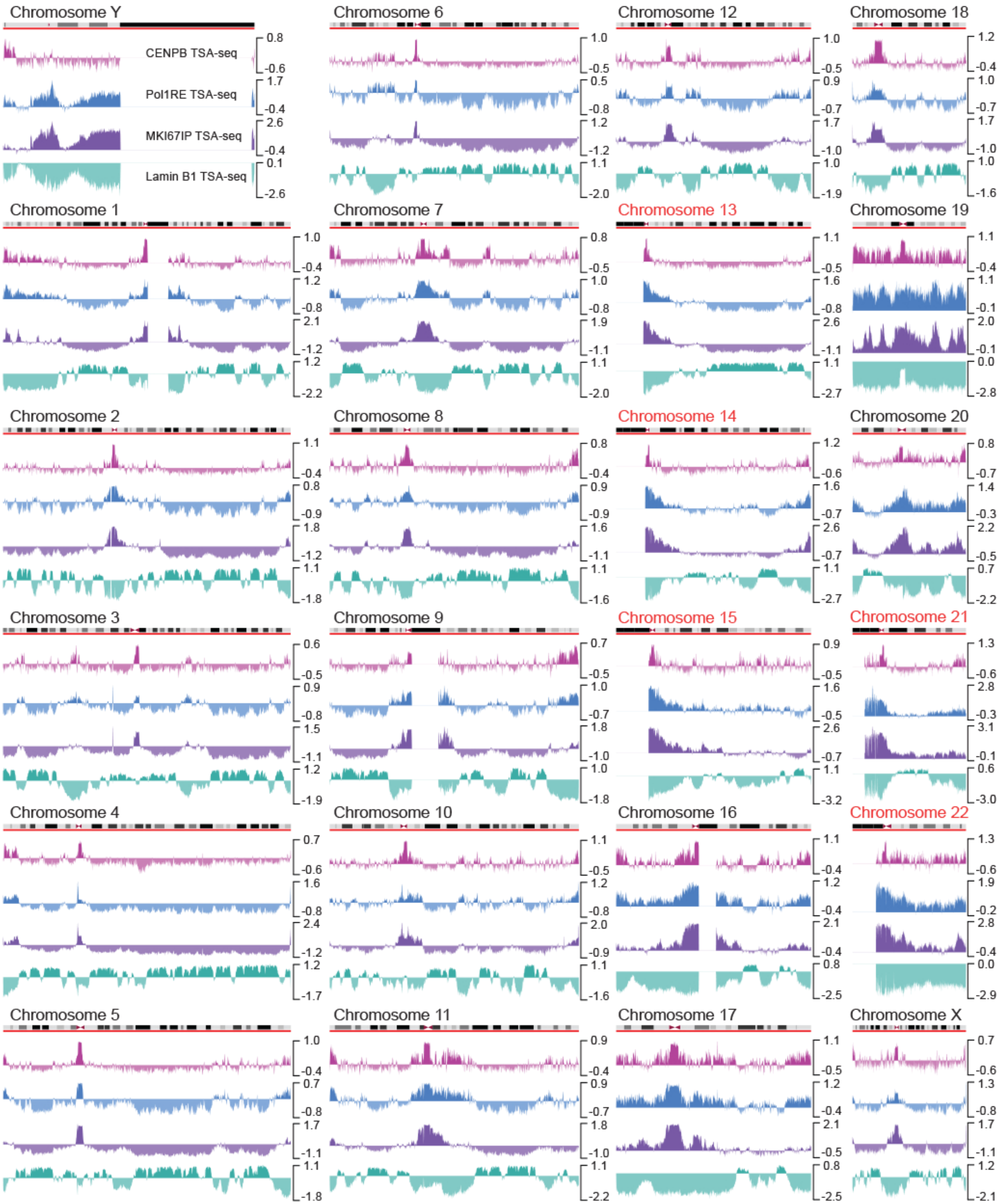

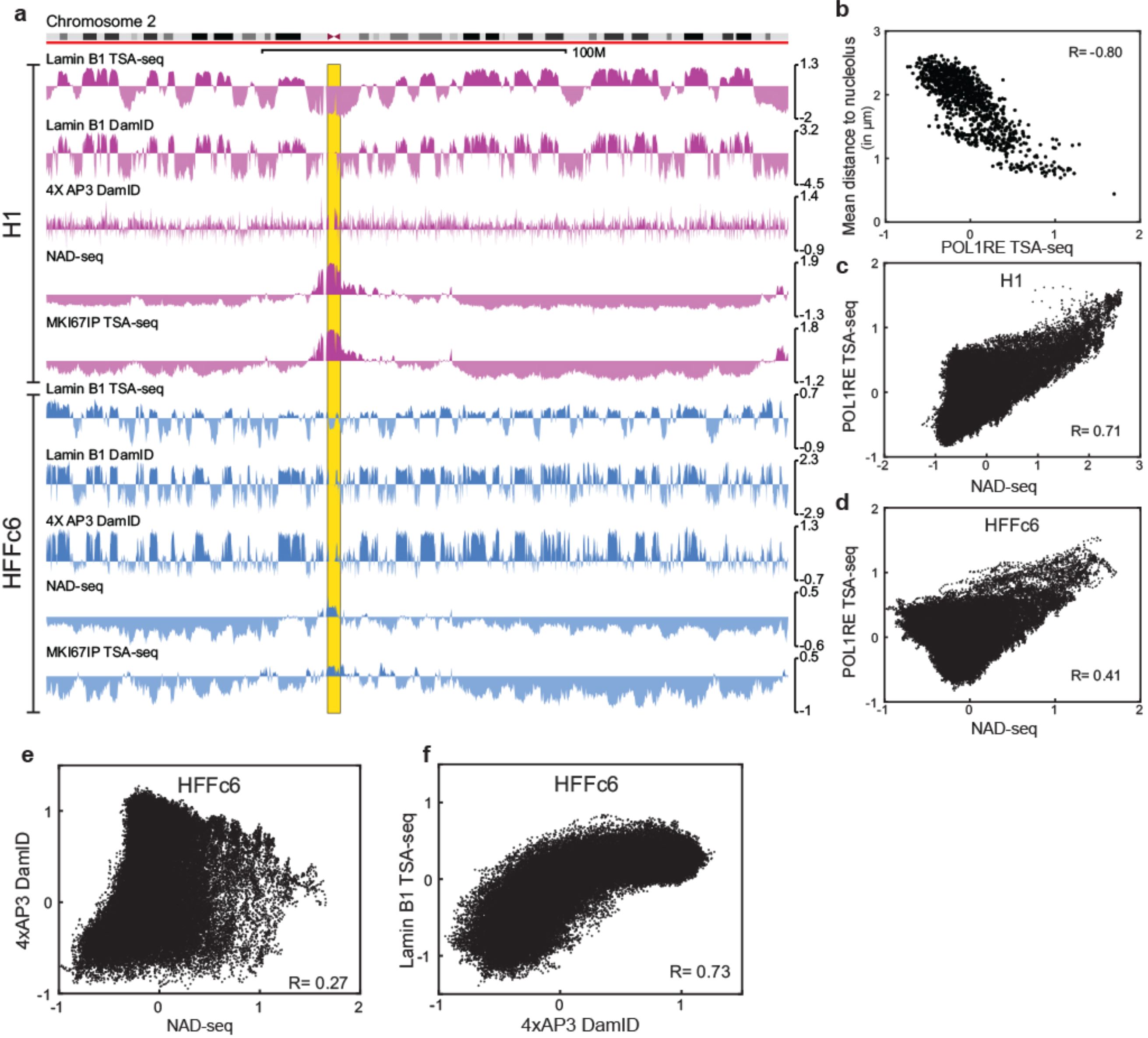

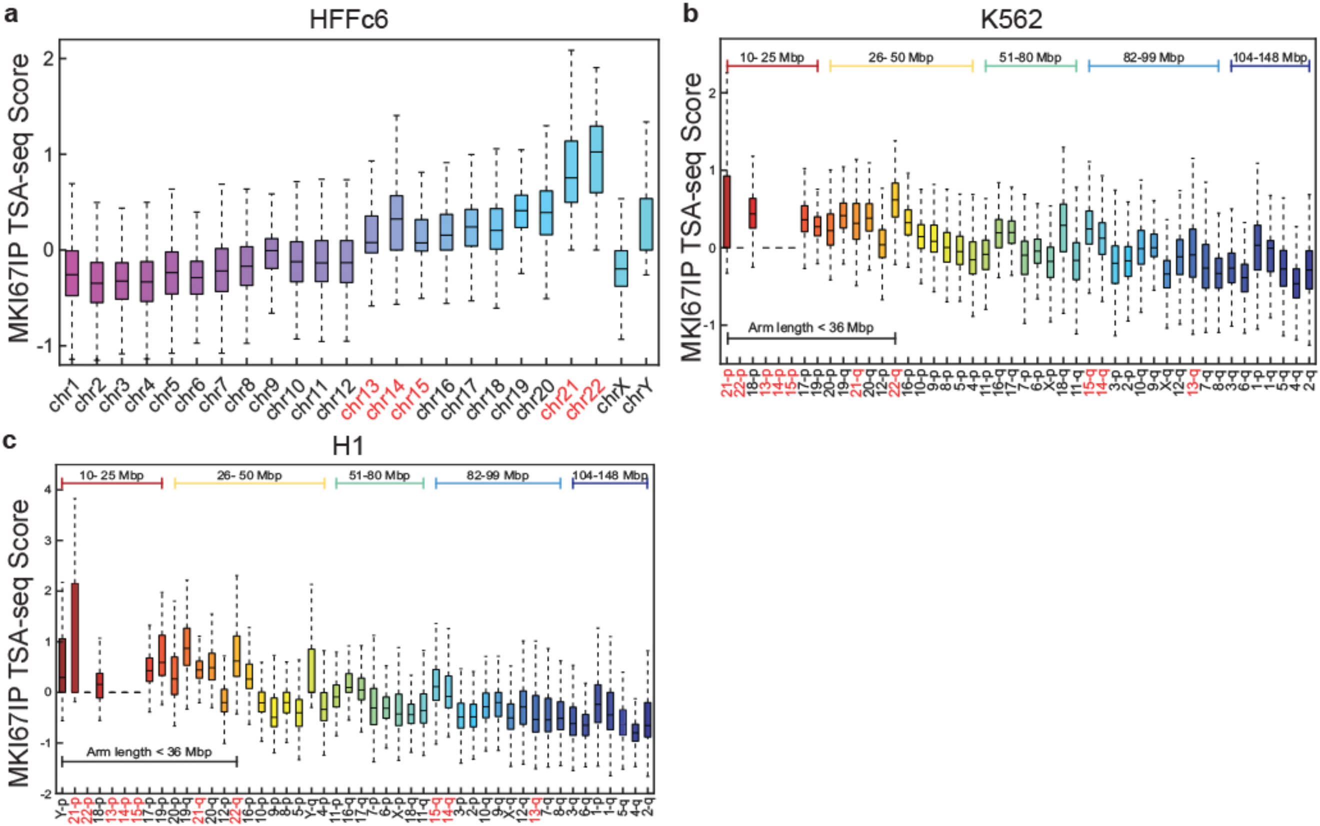

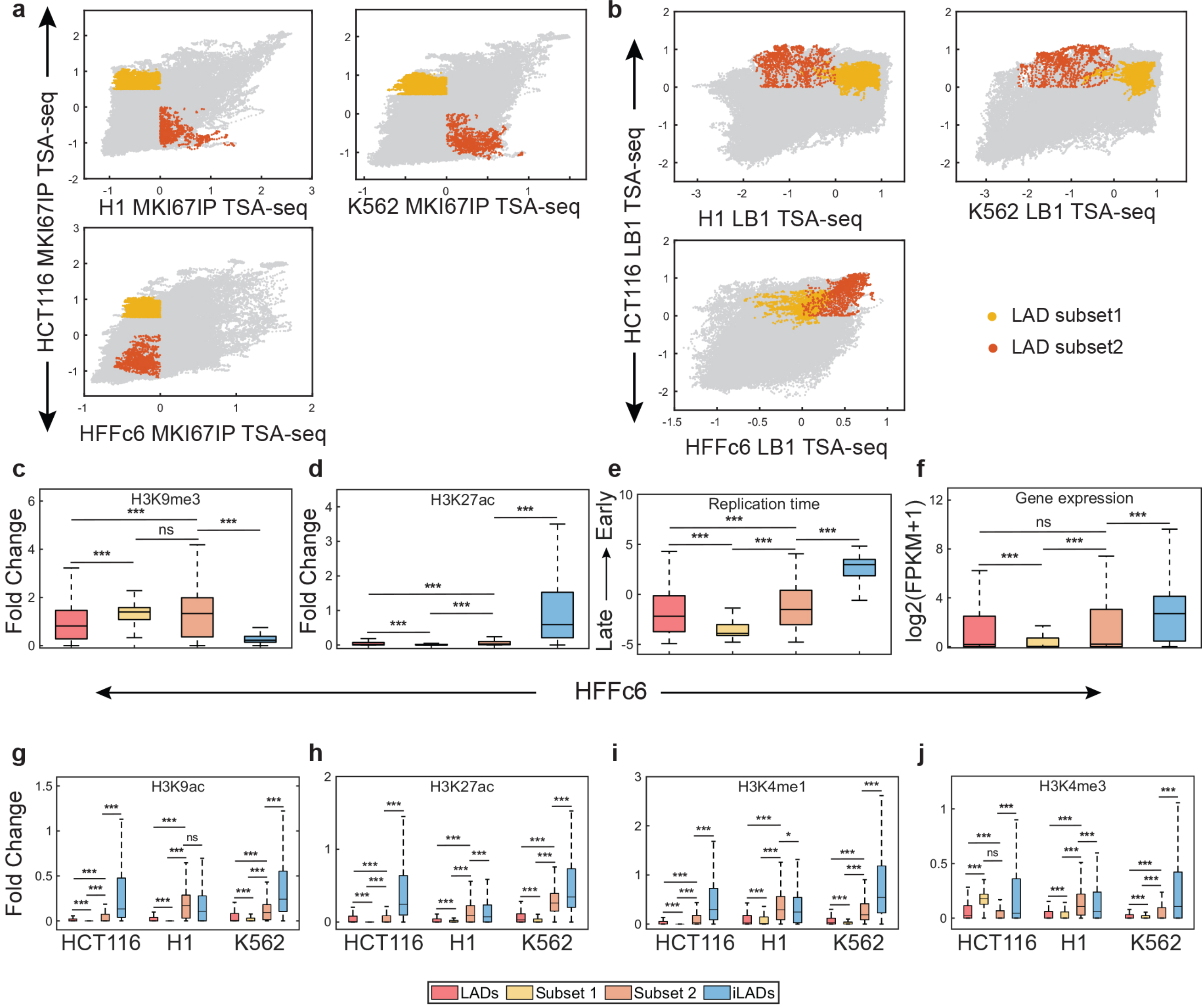

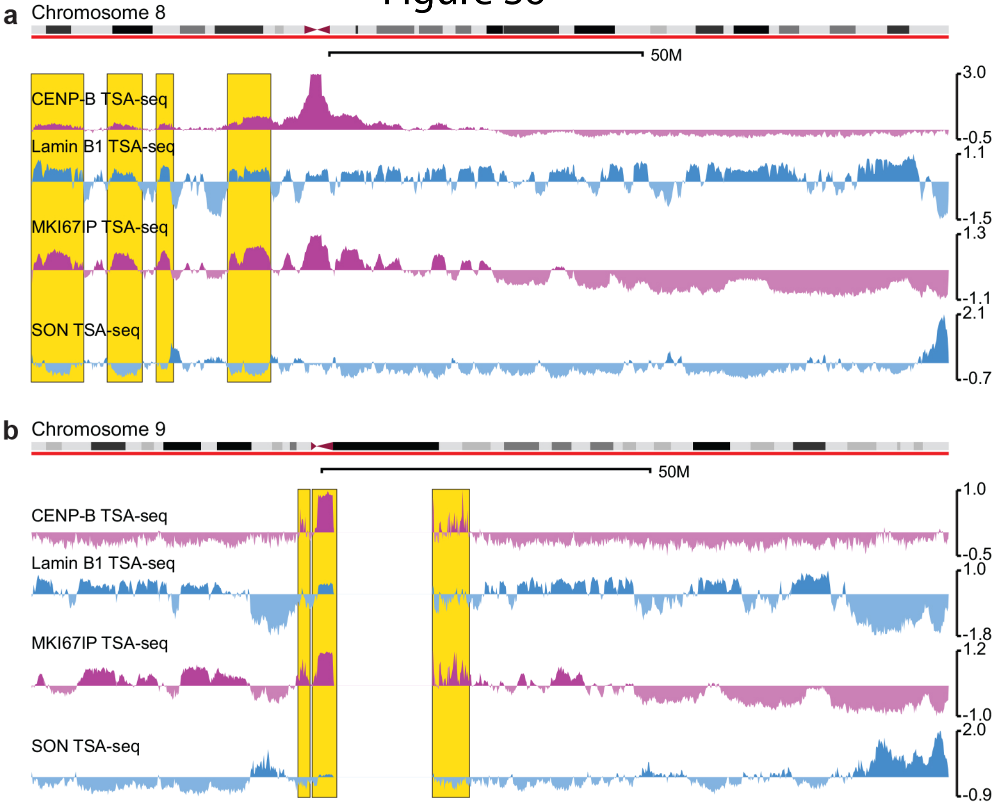

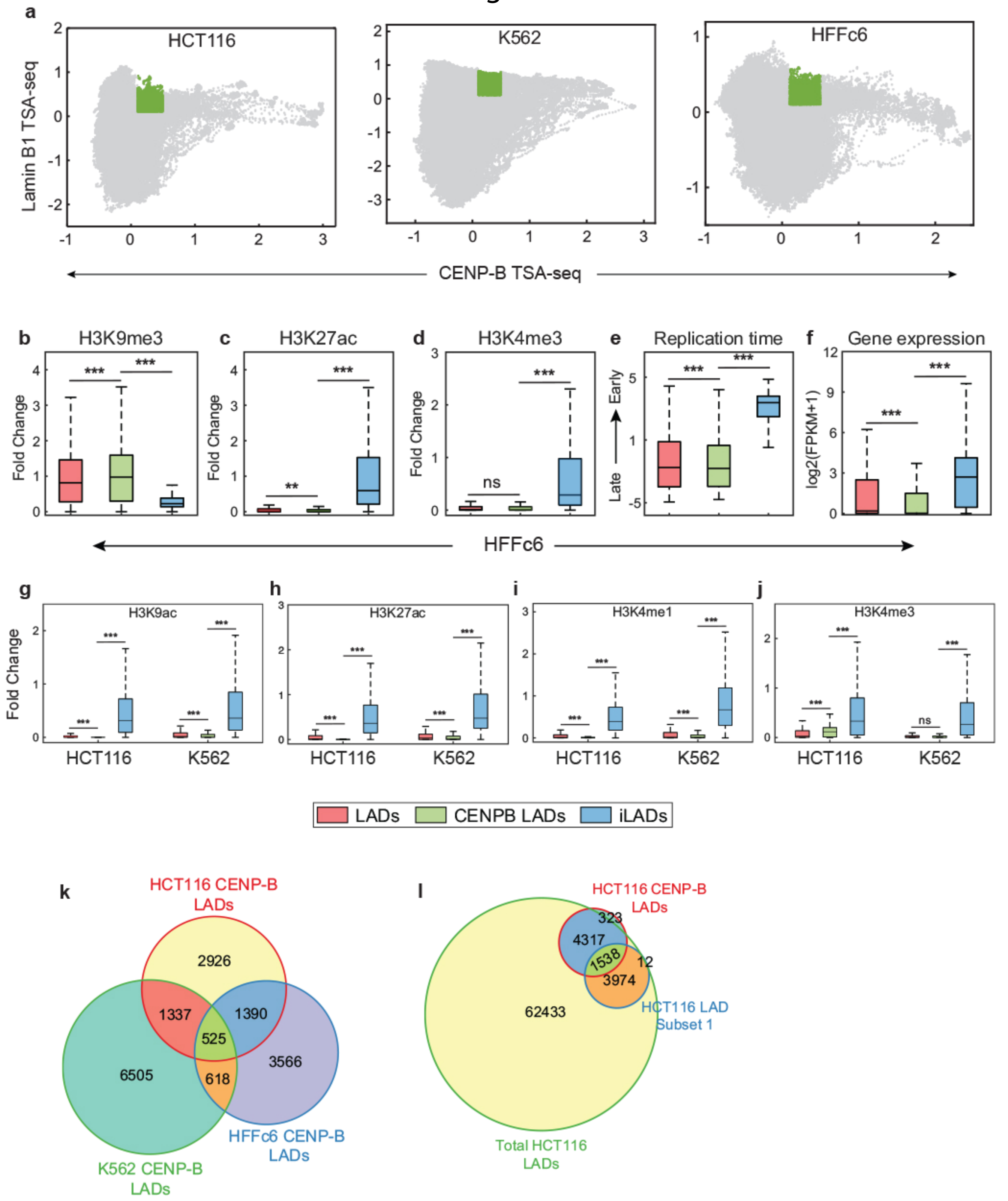

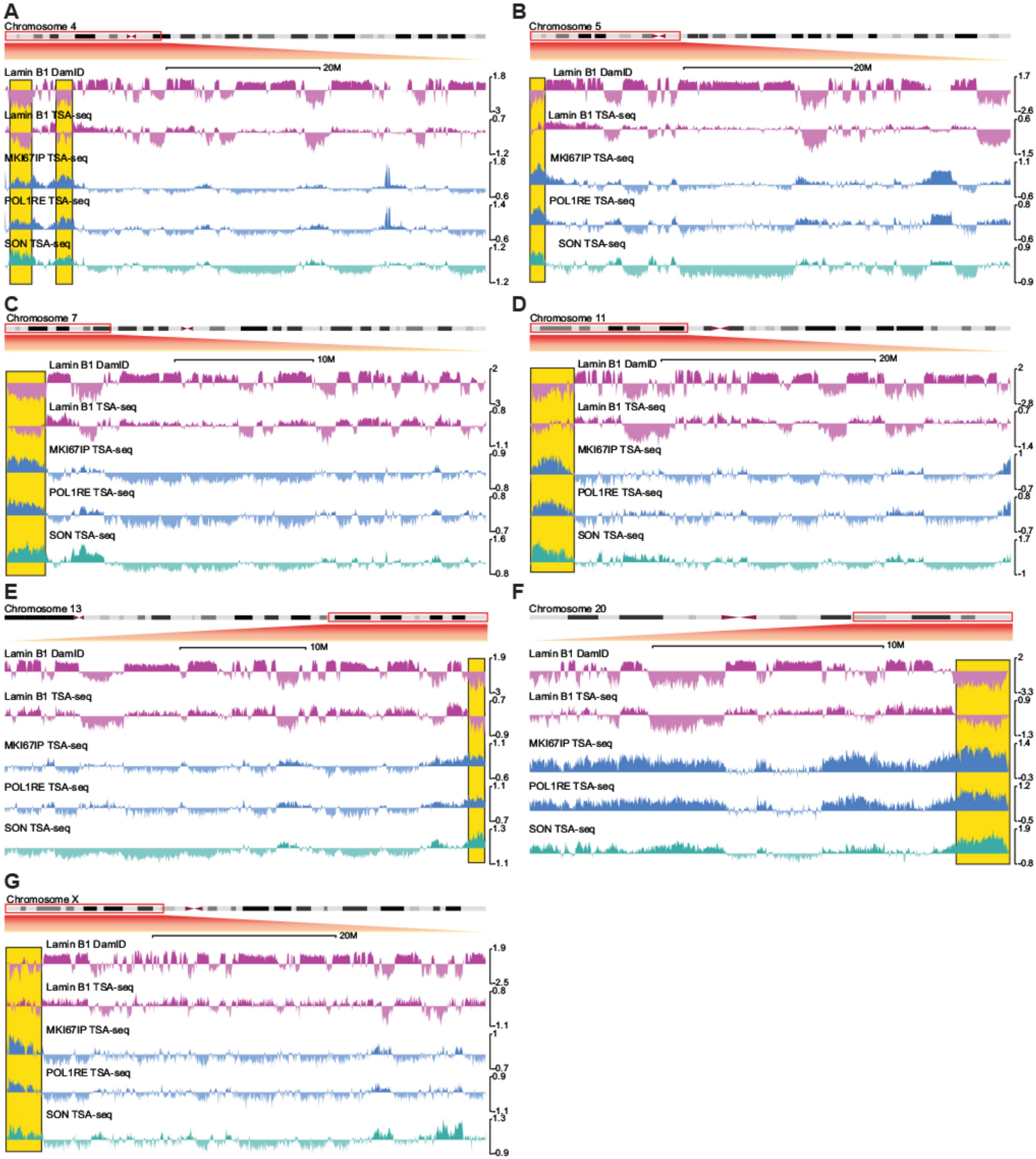
